# Benchmarking DNA Methylation Assays for Marine Invertebrates

**DOI:** 10.1101/2020.02.10.943092

**Authors:** Groves Dixon, Mikhail Matz

## Abstract

Interrogation of chromatin modifications, such as DNA methylation, has potential to improve forecasting and conservation of marine ecosystems. The standard method for assaying DNA methylation (Whole Genome Bisulfite Sequencing), however, is too costly to apply at the scales required for ecological research. Here we evaluate different methods for measuring DNA methylation for ecological epigenetics. We compare Whole Genome Bisulfite Sequencing (WGBS) with Methylated CpG Binding Domain Sequencing (MBD-seq), and a modified version of MethylRAD we term methylation-dependent Restriction site-Associated DNA sequencing (mdRAD). We evaluate these three assays in measuring variation in methylation across the genome, between genotypes, and between polyp types in the reef-building coral *Acropora millepora*. We find that all three assays measure absolute methylation levels similarly, with tight correlations for methylation of gene bodies (gbM), as well as exons and 1Kb windows. Correlations for differential gbM between genotypes were weaker, but still concurrent across assays. We detected little to no reproducible differences in gbM between polyp types. We conclude that MBD-seq and mdRAD are reliable cost-effective alternatives to WGBS. Moreover, the considerably lower sequencing effort required for mdRAD to produce comparable methylation estimates makes it particularly useful for ecological epigenetics.

## Introduction

The alarming effects of climate change on marine environments have led to a growing interest in Ecological Epigenetics. This relatively new field, focused on the interrelationships between environment, epigenetic modification, gene expression, and phenotypic variation (Bossdorf et al. 2008), has potential to improve forecasting and conservation of marine ecosystems. For instance, epigenetic modifications are hypothesized to mediate phenotypic plasticity, a mechanism important for resilience to environmental change (Reusch 2013; Eirin-Lopez and Putnam 2019). In humans, individuals prenatally exposed to famine show persistent differences in DNA methylation at relevant genes alongside alterations in disease risk (Painter et al. 2005; Heijmans et al. 2008). There is evidence that effects may extend even to the grandchildren of those who experienced food shortage (Kaati et al. 2007). Evidence from other mammals adds further support for such intergenerational, and even transgenerational effects (Radford et al. 2014; Irmler et al. 2020). In one remarkable case, traumatic olfactory conditioning in male mice was reported to produce epigenetic effects in F1s, and behavioral sensitivity even in F2s (Dias and Ressler 2014). Intergenerational effects and maternal effects have also been reported in plants (Feil and Fraga 2012), corals (Putnam and Gates 2015) and sea urchins (Wong et al. 2018; Strader et al. 2019; Wong et al. 2019). While such reports are exciting, it is important to maintain a reserved view on the overall importance of epigenetics for adaptation, especially as many published examples await independent replication (Horsthemke 2018) or have had attempts at replication fail to produce the same results (Irmler et al. 2020).

A notable feature found in plants and invertebrates is an association between gene body methylation (methylation of CpG sites within coding regions; gbM), and gene expression. In both groups, genes with gbM tend to be actively and stably expressed, whereas those without gbM tend toward less active, inducible expression (Zemach and Zilberman 2010; Sarda et al. 2012; Takuno and Gaut 2012; Gavery & Roberts, 2013; Takuno and Gaut 2013; Dixon et al. 2014; Dimond & Roberts, 2016; Takuno et al. 2016). Although gbM does not systematically regulate gene expression in plants or animals (Bewick et al. 2016; Zilberman 2017; Bewick et al. 2018; Bewick et al. 2019; Harris et al. 2019; Choi et al. 2020), comparisons between populations may still be ecologically informative. Indeed, in the coral *Acropora millepora*, comparative methylomics predicted fitness characteristics of transplanted corals better than either SNPs or gene expression (Dixon et al. 2018). The potential to predict fitness in novel conditions is especially important for conservation efforts involving outplanting individuals to maintain and rescue wild populations (van Oppen et al. 2015; van Oppen et al. 2017). Hence there is a need for cost-effective examination of chromatin modifications in ecological contexts. While chromatin marks such as histone modifications are undoubtedly important, DNA methylation is currently the easiest to measure, and the best-studied (Hofmann 2017).

Here, we use a model reef-building coral, *Acropora millepora*, to benchmark methods for assaying DNA methylation. Reef-building corals are prime candidates for the application of ecological epigenetics. They are exceptional both in their socio-ecological value, and sensitivity to anthropogenic change (Cesar 2000; Foden et al. 2013). As they are long-lived and sessile, they cannot migrate in response to suboptimal conditions, and must instead depend upon plasticity. Using this system, we compare three assays for measuring DNA methylation: Whole Genome Bisulfite Sequencing (WGBS), Methylated CpG Binding Domain Sequencing (MBD-seq)(Serre et al. 2009), and a modified version of the MethylRAD (Wang et al. 2015). WGBS, considered the gold standard for measuring DNA methylation, works by chemical conversion of unmethylated cytosines to uracils. Following PCR amplification, these bases are read as thymines. Hence, when mapped against a reference, fold coverage of reads indicating cytosine at a given site relative to fold coverage indicating thymines quantifies the rate at which the site was methylated in the original DNA isolation. MBD-seq works by capturing methylated DNA fragments with methyl-CpG-binding domains affixed to magnetic beads. This methodology has been used previously for ecological studies in *A. millepora* (Dixon et al. 2016; Dixon et al. 2018) and benchmarked against bisulfite sequencing in cultured embryonic stem cells (Harris et al. 2010). MethylRAD selects for methylated DNA through the activity of methylation-dependent restriction enzymes. DNA is digested with these enzymes, producing sticky ends exclusively near methylated recognition sites that allow for adapter ligation and sequencing. Methylation is quantified based on resulting fold coverage within a given region. The original MethylRAD protocol involved size selection for short fragments that were cut on both sides of palindromic methylated recognition sequences (Wang et al. 2015). We have modified the protocol by size-selecting for all digestion-derived fragments in the 170-700 bp range. The method is now conceptually similar to the genotyping by sequencing (GBS) protocol described in Elshire et al. (2011) and Andrews et al. (2016). To differentiate it from the original methylRAD, we refer to it as methylation-dependent Restriction site-Associated DNA sequencing (mdRAD).

With these three assays, we examine variation in methylation between genomic regions, between two polyp types (axial and radial), and between coral colonies (genotypes). We compare results from each assay to assess how consistently they measure methylation, and the optimal sequencing effort to maximize sensitivity while minimizing costs.

## Materials and Methods

### Sample collection

Two adult colonies of *A. millepora* were collected by SCUBA on November 25^th^, 2018, one from Northeast Orpheus (labeled N12), and one from Little Pioneer Bay (labeled L5), under the Great Barrier Reef Marine Park Authority permit G18/41245.1. Colonies were maintained in the same raceway with flow of unfiltered seawater for 22 days. Branches from each colony were submerged in 100% ethanol and immediately placed at −80°C for 48 hours. Samples were then maintained at −20 or on ice for approximately 48 hours during transport to the laboratory where they were again stored at −80°C until processing.

### DNA Isolation

For each axial polyp sample, the very tips of four branches were cut off and pooled. For radial polyps, similar amounts of tissue were pooled from the sides of the same four branches. Tissue samples were lysed in Petri dishes with 2 ml of lysis buffer from an RNAqueous™ Total RNA Isolation Kit (cat no. AM1912). DNA was isolated using phenol:chloroform:isoamyl alcohol with additional purification using a Zymo DNA cleanup and concentrator kit (cat no. D4011)(Supplemental Methods file). Isolations were quantified using a Quant-iT™ PicoGreen™ dsDNA Assay Kit (cat no. P7589). The same DNA isolations were used for each downstream methylation assay. We isolated three replicates from each genotype-tissue pairing, for a total of 12 isolations (2 colonies, 2 tissues, 3 replicates per). In downstream analyses, we use treatment groups to refer to either coral colony (N12 vs L5), or polyp type (tip vs side).

### Whole genome bisulfite sequencing library preparation

Whole genome bisulfite sequencing (WGBS) libraries were prepared using a Zymo Pico Methyl-Seq Library Prep Kit (cat no. D5455). Each library was prepared from 100 ng of genomic DNA. For half the samples, we included 0.05 ng (0.05%) of λ phage standard DNA to estimate conversion efficiency. The final sample size was 8 (2 genotypes, 2 tissues, 2 replicates per; Table 1). The 8 libraries were sequenced across four lanes on a Hiseq 2500 for single-end 50 bp reads at The University of Texas Austin Genome Sequencing and Analysis Facility (GSAF). Single-end sequencing was recommended in the Zymo Pico Methyl-Seq manual.

**Table 1.** Sample and library information.

| Assay | Treatment groups | Replicates | samples | Library types | Total Libraries | Raw reads | Final aligned reads |
| --- | --- | --- | --- | --- | --- | --- | --- |
| WGBS<sup>1</sup> | 4 | 2 | 8 | 1 | 8 | 9.54E+08 | 3.73E+08 |
| MBD-seq<sup>2</sup> | 4 | 2 | 8 | 2* | 16 | 4.88E+08 | 3.94E+08 |
| mdRAD | 4 | 3 | 12 | 2** | 24 | 2.85E+08 | 1.23E+08 |

<sup>1</sup>Zymo Picomethyl Kit
<sup>2</sup>Diagenode Methylcap Kit
\*Both captured and unbound fractions were sequenced
\*\*Separate libraries prepared with Fspe1 and Mspj1

### mdRAD library preparation

mdRAD libraries were prepared using a protocol based on Wang et al. (2015). Importantly, Wang et al. (2015) selected small sized fragments that had been cut on either end by the enzyme due to palindromic recognition sequences. Since in our hands the yield of the palindrome-derived product was very low, we instead sequenced any ligated fragments in the 170-700 bp range. We also used different oligonucleotide sequences, designed for similarity to those used in the current 2bRAD protocol (Table S1)(Dixon et al. 2015; Matz et al. 2018; Matz 2019). A detailed version of the protocol used is included as a Supplemental Methods file. We prepared libraries using two different methylation-dependent endonucleases, FspE1 (NEB cat no. R0662S) and MspJ1 (NEB cat no. R0661S). For each library, we used 100 ng of genomic DNA as input. Digests were prepared with 0.4 units of endonuclease and the recommended amounts of enzyme activator solution and Cutsmart buffer (final volume = 15.0 µl) and incubated at 37°C for four hours. We then heated the digests to deactivate the enzymes for 20 minutes (at 80°C for FspE1 and 65°C for MspJ1). All ligations were prepared with 0.2 µM mdRAD 5ILL adapter, 0.2 µM of the mdRAD 3ILLBC1 adapter, 800 units of T4 ligase, 1mM ATP (included in ligase buffer), and 10 µl of digested DNA (final volume = 20 µl). Ligations were incubated at 4°C overnight (approximately 12 hours). Ligase was then heat-inactivated by incubation at 65°C for 30 minutes. Sequencing adapters and multiplex barcodes were then appended by PCR. Each PCR was prepared with 0.3 mM each dNTP, 0.15 µM of the appropriate ILL_Un primer, 0.15 µM of the appropriate ILL_BC primer, 0.2 µM of the p5 primer, 0.2 µM of the p7 primer, 1x Titantium taq buffer, 1x Titantium taq polymerase, and 7 µl of ligation (final volume = 20 µl)(Table S1). At this point in the protocol, all samples were distinguishable by the dual barcoding scheme. The concentration of each PCR product was quantified using PicoGreen™ dsDNA Assay Kit (cat no. P7589). Based on these concentrations, 200 ng of each product was combined into a final pool with approximate concentration of 32 ng/µl. A portion of this pool was then size selected for 170 – 700 bp fragments using 2% agarose gel and purified using a QIAquick gel Extraction kit (cat no. 28704). After gel purification, the pool was sequenced with a single run on a NextSeq 500 for paired-end 75 bp reads at the University of Texas Genome Sequencing and Analysis Facility. The final number of libraries included in the pool was 24 (2 genotypes, 2 tissues, 2 different restriction endonucleases, 3 replicates per combination; Table 1). As this methylation assay depends on fold coverage to infer methylation levels, single-end reads are a more cost-effective approach. We opted for paired-end reads in this case only to ensure proper product structure for benchmarking purposes.

### MBD-seq library preparation

MBD-seq libraries were prepared using a Diagenode MethylCap kit (cat no. C02020010) as described previously (Dixon et al. 2016; Dixon et al. 2018). Briefly, genomic DNA was sheared to a target size of 300 – 500 bp. Concentrations based on PicoGreen™ dsDNA assay on genomic DNA were assumed not to have changed during shearing. Because limited genomic DNA remained, we prepared these libraries from pools of genomic DNA for each genotype-tissue pair. Also due to limited genomic DNA, the two libraries for N12 tips were prepared using only 0.565 μg as input. For the remaining libraries, half were prepared with 1 μg of input and the other half from 1.5 μg. During capture of methylated DNA, we retained the flow-through for sequencing, which we refer to at the unbound fraction. Captured methylated fragments were eluted from capture beads in one single total elution using High Elution Buffer. The final sample size was 8 (2 genotypes, 2 tissues, 2 replicates per; Table 1). After capture, fragment size was assessed using 1.5% agarose gels. The captured and unbound fractions both ranged between 200 and 1000 bp. These fragments were submitted to the University of Texas Genome Sequencing and Analysis Facility. Here the fragments were further sheared to a target size of 400 bp. This additional shearing was done to ensure appropriate library sizes of 300 – 500 bp for sequencing. Libraries were prepared with a NEBNext Ultra II DNA Library Preparation Kit (cat no. E7645). Libraries were sequenced with a single run on a NextSeq 500 for single-end 75 bp reads.

### Whole genome bisulfite sequencing data processing

Raw reads were trimmed and quality filtered using cutadapt, simultaneously trimming low-quality bases from the 3’ end (-q 20) and removing reads below 30 bp in length (-m 30)(Martin 2011). Trimmed reads were mapped to the *A. millepora* reference genome (Fuller et al. 2019) using Bismark v0.17.0 (Krueger and Andrews 2011) with adjusted mapping parameters (--score_min L,0,-0.6) in --non_directional mode as indicated in the Pico Methyl-Seq Library Prep Kit manual. Methylation levels were extracted from the alignments using bismark_methylation_extractor with the --merge_non_CpG, -- comprehensive, and --cytosine_report arguments. At this point, CpG sites within the lambda DNA chromosome and the mitochondrial chromosome were set aside to assess conversion efficiency. Conversion efficiencies were estimated as the ratio of ‘unmethylated’ fold coverage (converted by bisulfite treatment) to all fold coverage summed across CpG sites in the lambda DNA and the host mitochondrial reference sequences. Detailed steps used to process the WGBS reads are available on the git repository (Dixon 2020).

### MBD-seq data processing

Raw reads were trimmed and quality filtered using cutadapt simultaneously trimming low-quality bases from the 3’ end (-q 20) and removing reads below 30 bp in length (-m 30)(Martin 2011). Trimmed reads were mapped to the *A. millepora* reference genome (Fuller et al. 2019) with bowtie2 using the --local argument (Langmead and Salzberg 2012). Alignments were sorted and indexed using samtools (Li et al. 2009), and PCR duplicates were removed using MarkDuplicates from Picard Toolkit (Broad Institute 2019). Fold coverage for different regions (eg. gene boundaries, exon boundaries, 1 Kb windows, etc.) was counted using multicov from BEDTools (Quinlan and Hall 2010). Detailed steps used to process the MBD-seq reads are available on the git repository (Dixon 2020).

### mdRAD data processing

*A*ll mdRAD reads were expected to contain NNRWCC as the first six bases of the forward read, and ACAC as the first four bases of the reverse read (Table S1; Supplementary Methods Section). The degenerate NNRW sequence in the forward read allows for discrimination of PCR duplicates, as uniquely ligated digestion products are unlikely (1/64) to bear identical sequences for these four bases. With this in mind, we used a custom python script to filter out any reads for which the first 20 bp was duplicated in a previous read (ie a likely PCR duplicate). At the same time, all paired end reads were filtered to retain only those with the expected NNRWCC beginning to the forward read and ACAC in the reverse read. These non-template bases were trimmed, along with adapters and low-quality bases using cutadapt (Martin 2011). Trimmed reads were mapped to the *A. millepora* reference genome (Fuller et al. 2019) with bowtie2 using the --local argument (Langmead and Salzberg 2012). Alignments were sorted and indexed using samtools (Li et al. 2009). Fold coverage for different was counted using multicov from BEDTools (Quinlan and Hall 2010). Detailed steps used to process the mdRAD reads are available on the git repository (Dixon 2020).

#### Designating of regions of interest

Statistical analyses for all three assays were based on windows recorded in .bed files. These included genes, exons, upstream sequences, and tiled windows of varying sizes. Gene, exon, and upstream sequence boundaries were identified from the reference GFF file (Fuller et al. 2019). Upstream sequences included 1 Kb upstream of each gene. These were intended to approximate promoter regions. Tiled windows were generated using makewindows from the BEDTools suite (Quinlan and Hall 2010). General statistics for these regions such as length, nucleotide content, and the number of CpGs, were extracted from the reference genome with a custom python script using SeqIO from Biopython (Cock et al. 2009). All downstream analyses of methylation level and differences between groups were based on these regions.

### Whole genome bisulfite statistical analysis

Statistical analyses of WGBS data were conducted on the .cov files output from Bismark. Analysis was conducted only on CpG sites. Methylation level was calculated in several ways. The simplest metric was the overall fractional methylation, calculated as the number of methylated counts divided by all counts summed across CpGs within the region.

We report this as the % methylation on the log_2_ scale throughout the manuscript (eg Figure 1A). We calculated a similar metric using generalized logistic regression. Here the estimate of a region’s methylation level was the sum of the intercept and the region’s coefficient for a model of the probability of methylation given all methylated and unmethylated counts within the region. We also report the frequency of methylated CpGs, calculated as the number of methylated CpG sites divided by the total number of CpG sites within a region. We classified a CpG as methylated when the number of methylated counts was significantly greater than the null expectation with 0.01 error rate (binomial test; p-value < 0.05). We also calculated the ratio of methylated CpGs to the total length (bp).

**Figure 1:**
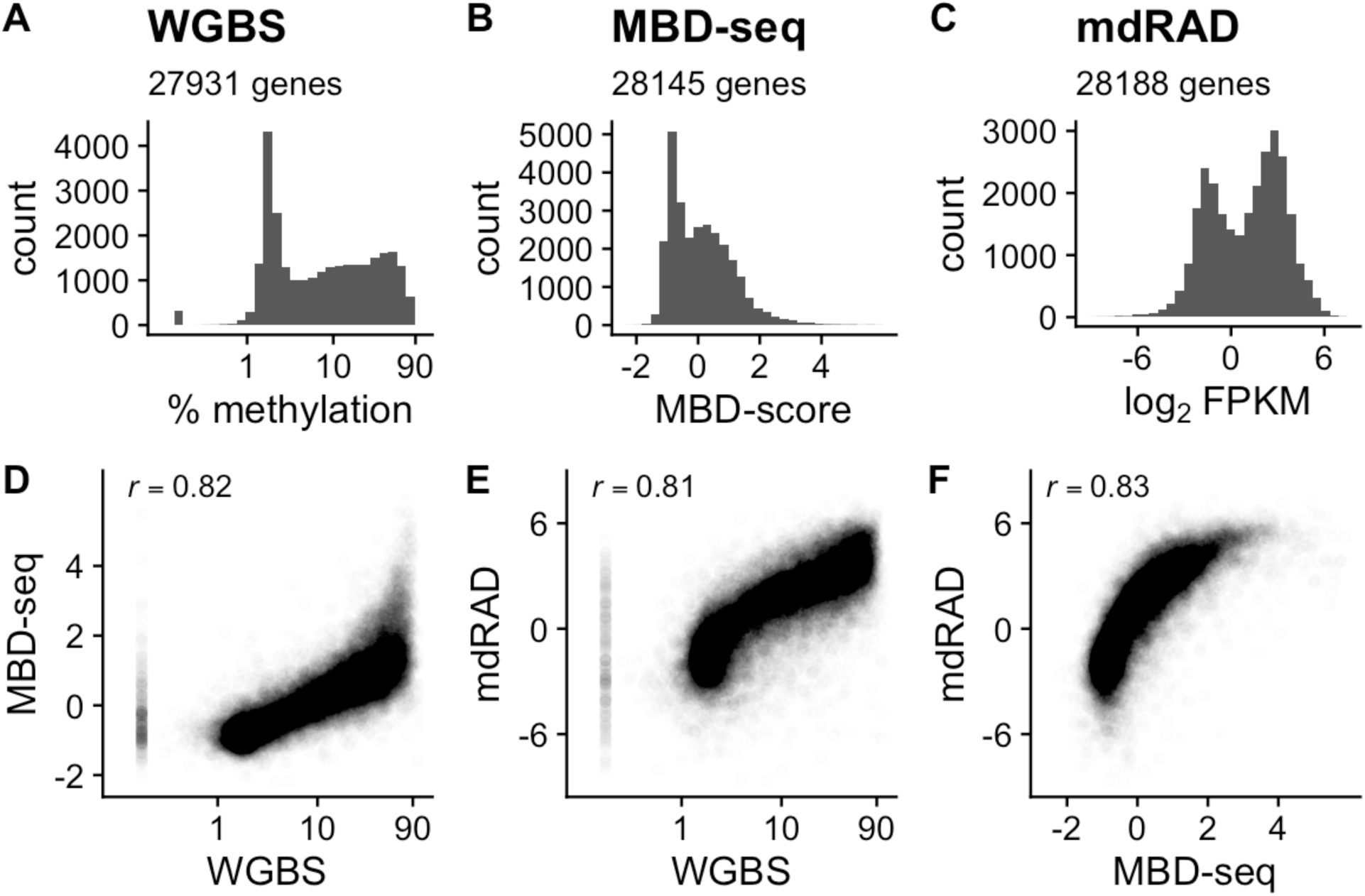
Correlation of gbM level estimates from each assay. (A-C) Histograms of gbM level. (A) WGBS. Axis is on the log scale. (B) MBD-seq. MBD-score refers to the log2 fold difference between the captured (methylated) and unbound (unmethylated) fractions from the library preparation. (C) mdRAD. Plot shows log_2_ FPKM from combined reads from both enzymes. (D-E) Scatterplots of methylation level estimates from each assay. Pearson correlations are indicated in the top left.

Statistical analysis of differences in methylation between treatment groups (tissue type or colony) was done with the MethylKit package (Akalin et al. 2012). Filtering parameters supplied to the filterByCoverage() function were lo.count=5, and hi.perc=99.9. The function methylKit::unite() was run using min.per.group = 4, so that only sites with data from all samples in each treatment group passed. Methylation counts for particular regions were isolated using the appropriate .bed file, the Granges() function from the GenomicRanges package (Lawrence et al. 2013), and the regionCounts() function from MethylKit.

### MBD-seq statistical analysis

Statistical analyses of MBD-seq data were conducted on the fold coverages output from BEDTools multicov. Methylation level was calculated based on the difference in fold coverage between the captured and unbound fractions taken during library preparation. We quantified this using DESeq2 as the log_2_ fold change between the two fractions from a model including colony and polyp type as covariates (Love et al. 2014). Following previous studies (Dixon et al. 2016; Dixon et al. 2018), we refer to this value as the MBD-score. We also calculated methylation level based on the fragments per kilobase per million reads (FPKM) from the captured fraction averaged across all samples. Differential methylation was also assessed using DESeq2. This was done in two ways, one using both the captured and unbound fractions, the other using only the captured fraction. Using both the captured and unbound fractions, the effect of treatment group was assessed as the interaction between treatment group and fraction. In other words, we assessed the effect of treatment group on the difference between the captured and unbound fractions. To assess methylation differences without using the unbound fraction, we simply compared fold coverages from the captured fraction between treatment groups with a model including the alternative grouping as a covariate. DESeq tests were run using fitType = ‘local’ and significance was assessed using Wald tests.

### mdRAD statistical analysis

Statistical analyses of mdRAD data were conducted on the fold coverages output from BEDTools multicov. Methylation level was calculated as FPKM averaged across all samples, and as the fragments per recognition site per million reads. Methylation differences were calculated using DESeq2 comparing fold coverage between treatment groups while controlling for the restriction enzyme used and the other treatment group. DESeq tests were run using fitType = ‘local’ and significance was assessed using Wald tests.

### Simulating reduced fold coverage

To assess the importance of fold coverage for methylation statistics, we simulated reduced fold coverages for each of the three assays. For MBD-seq and mdRAD, this was done by sampling iteratively lower total counts with replacement weighted by the gene’s proportion of total read counts in the original dataset. To clarify, to simulate read reductions for 28188 genes for each sample, a vector of weights was generated by dividing each gene’s fold coverage by the total for the sample. A vector ranging from 1 to 28188 was then randomly sampled with replacement, with probabilities set by the weight vector. The number of times each value was sampled was then totaled to give each genes’ count in the simulated fold reduction. For WGBS, the trimmed fastq files were randomly sampled without replacement and all processing steps were repeated as indicated above.

### Statistical reporting

Unless otherwise noted, we report significant results as those with false discovery corrected p-values less than 0.1 (FDR < 0.1)(Benjamini and Hochberg 1995). Correlations are reported as Pearson correlations. All scripts for data processing and analysis in this study are available on GitHub: (Dixon 2020).

## Results

### WGBS sequencing results

Sequencing the WGBS libraries produced 954 million single-end reads across 8 samples (2 from each colony-tissue type pair; median = 120 million per sample). Trimming and quality filtering reduced the median to 119 million per sample. Mapping efficiency was 40% on average, with a median of 47 million mapped reads per sample. Conversion efficiency averaged 98.5 ± se 0.05% based on spiked in lambda DNA and 98.0 ± se 0.10% based on mitochondrial DNA. The overall percentage of mapped reads was 39% of raw reads.

### MBD-seq sequencing results

Sequencing the MBD-seq libraries produced a total of 488 million single-end reads. These were divided across 8 samples each with two libraries (one captured and one unbound). Median read count for the captured and unbound libraries was 27.4 and 33.1 million respectively. Trimming and quality filtering removed 0.1% of reads. Mapping efficiency was 92% on average, with medians of 24.9 and 30.8 million reads for captured and unbound libraries respectively. PCR duplication rate was 12% on average, for final medians of 21.8 and 27.2 million mapped reads per sample for the captured and unbound fractions respectively. The final percentage of countable reads (passing all filters and properly mapped) was 78% of raw reads for captured libraries and 82% for unbound libraries.

### mdRAD sequencing results

Sequencing the mdRAD libraries produced a total of 284 million paired-end reads across 24 libraries (3 replicates for each of the 4 colony-polyp type combinations each prepared with 2 different restriction enzymes). These were filtered to include only reads with the appropriate adapter sequences found in both the forward and reverse directions (∼71% of reads) and to remove PCR duplicates based on degenerate sequences incorporated into the forward read (average 13.5% duplication rate). On average 60% of raw reads passed both these filters (172 million total passing reads). Trimming and quality filtering further reduced this by 0.2%, for 74 million reads for Fspe1 libraries (median = 5.9 million per sample) and 98.5 million for the Mspj1 libraries (median = 7.3 million per sample). Properly paired mapping efficiency averaged 77% and 66% for Fspe1 and Mspj1 libraries respectively, giving final median read counts of 4.6 and 4.9 million reads per library. The final percentage of raw reads that passed all filters and properly mapped was thus 44% for Fspe1 and 42% for MspJ1.

### Estimating methylation level

Measurements of absolute levels of gbM were consistent across assays. Each assay identified a bimodal distribution of gbM (Figure 1 A-C). Pearson correlations between assays were all greater than 0.8 (Figure 1 D-F). All three assays correlated negatively with the CpGo/e, with the strongest correlation for WGBS (Figure S1). Correlations similar to those for gbM were found for exons (Figure S2), 1 Kb windows (Figure S3), and upstream regions of coding sequences (1 Kb upstream from the gene boundary) (Figure S4).

The measures of gbM level shown in Figure 1 A-C were selected based on their simplicity and correlation between assays. Additional metrics of gbM level for WGBS, MBD-seq, and mdRAD are shown in figures (Figure S5; Figure S6; Figure S7). For WGBS, these included estimates based on logistic regression, the ratio of methylated CpGs to all CpGs, and ratio of methylated CpGs to gene length. Of these, all except ratio of methylated CpGs to gene length correlated roughly equivalently with the other two assays (Figure S5). For MBD-seq, metrics that did not include the unbound fraction (FPKM and a similar metric based on the number of CpGs) correlated poorly with other assays (Figure S6). Hence sequencing the unbound fraction is important for measuring absolute methylation level with MBD-seq. For mdRAD, the two restriction enzymes produced nearly equivalent results. mdRAD FPKM was more consistent with other assays than a similar metric based on the number of recognition sites (Figure S7).

### Methylation differences between groups

Estimates of differential methylation between coral colonies were concordant between assays, but less so than methylation level. Each assay identified extensive differential methylation between the two colonies (Figure 2A-C). The number of significant differentially methylated genes (DMGs) detected with each assay reflected the sample sizes used, rather than overall sequencing effort (Table 1). mdRAD, with 24 libraries, identified the most, with 12,464 DMGs. MBD-seq, with 8 pairs of captured and flow-through libraries, identified the second most (6,347 DMGs). WGBS, with 8 libraries, detected 4,395 DMGs. The overlap between these sets of DMGs is shown in Figure 3. Although it only used roughly 1/10^th^ of the sequencing effort, a reduced mdRAD dataset using only 8 libraries generated with FspE1 still identified 7407 DMGs (Figure S8).

**Figure 2:**
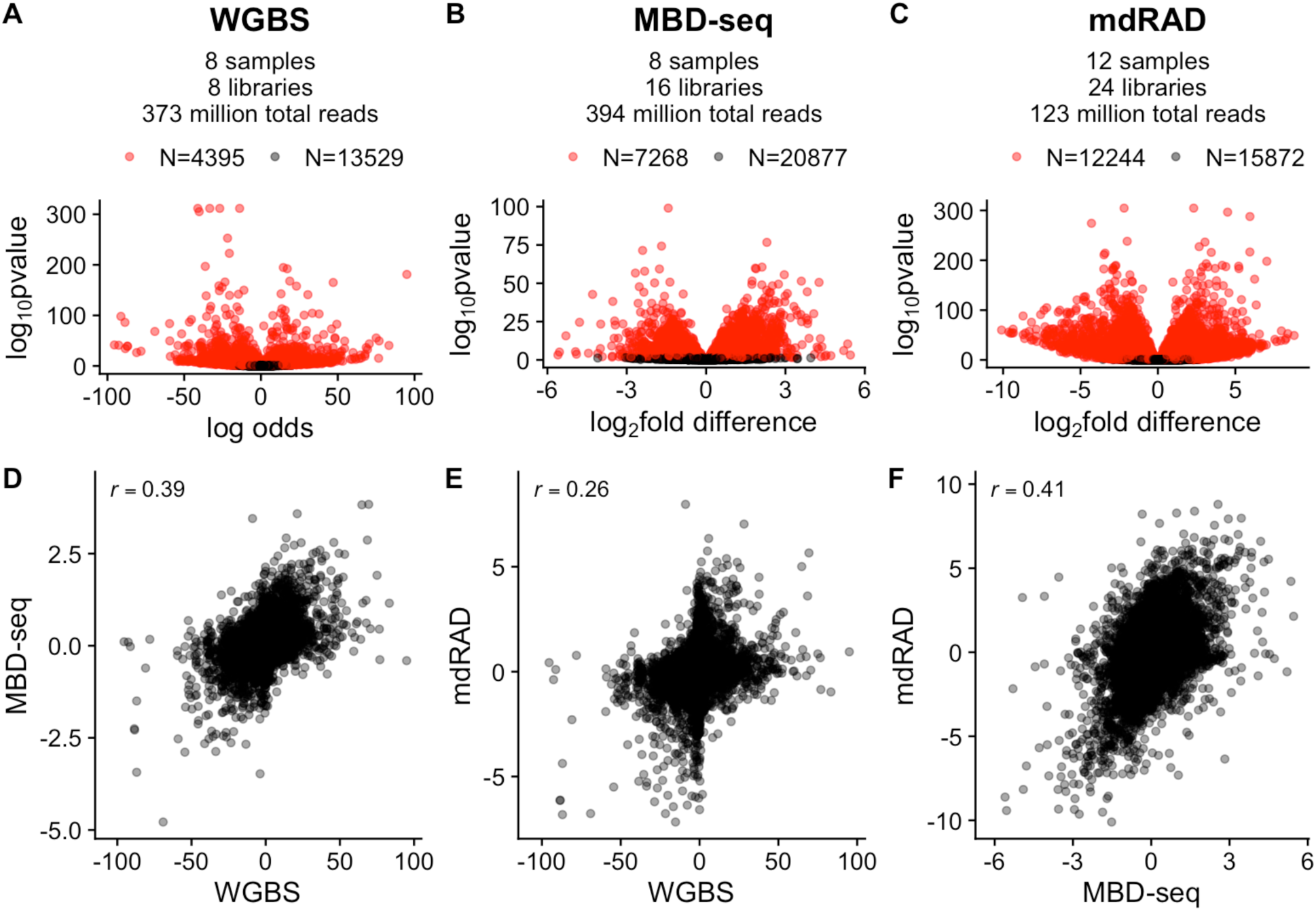
Correlation of gbM difference estimates between two coral colonies (genotypes). (A-C) Volcano plots illustrating differential gbM for the indicated assay. Red points indicate significant genes (FDR < 0.1). The number of biological samples, libraries, total number of filtered and aligned reads, and the number of significant and nonsignificant genes is given in the subtitle for each panel. (D-F) Scatterplots of gbM difference estimates for the indicated assays. Pearson correlations are indicated in the top left.

**Figure 3:**
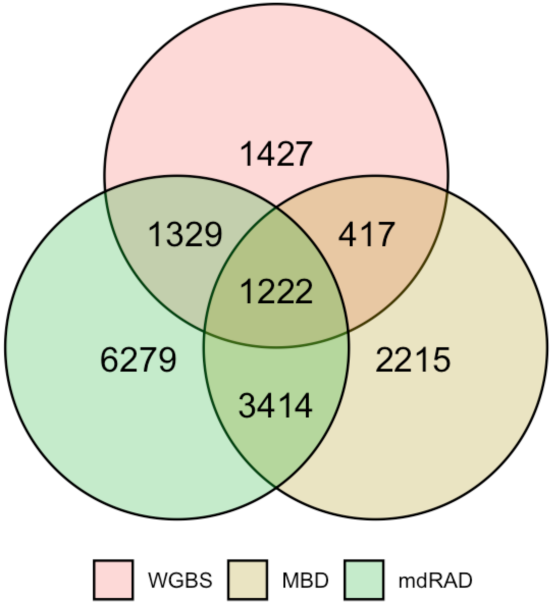
Venn diagram showing overlap of differentially methylated genes detected with each assay.

Despite variations between assays and statistical methods, estimates of methylation differences were positively correlated (Figure 2 D-F). MBD-seq correlated with the other two assays similarly (Pearson correlation = 0.39 and 0.41). mdRAD and WGBS were less correlated (Pearson correlation = 0.26). Correlations were stronger (0.31 – 0.55) when only methylated genes (> 3.1% methylation based on WGBS; Figure 1A) were considered. Similar results were found for differences between exons (Figure S9), 1 Kb windows (Figure S10), and upstream regions of coding sequences (Figure S11). Hence, estimates of methylation differences between colonies (genotypes) were noisy, but reproducible across assays.

In contrast to differential methylation between colonies, differences between polyp types were weak, and not reproducible across assays. The number of significant differences was reversed compared to the colony comparison, with the most (169 DMGs) detected by WGBS, the second (12 DMGs) by MBD-seq, and the least (1 DMG) by mdRAD (Figure S12). There was no overlap in significant calls between assays. Difference estimates based on WGBS showed no correlation with the other two assays (Pearson correlation between 0.01 and 0.02). MBD-seq and mdRAD correlated weakly 0.2 (Figure S12).

### Spatial precision

Correlations between assays were generally robust across window sizes. For each assay, we calculated methylation level, as well as methylation differences between the two colonies for tiled windows of varying sizes: (100bp, 500bp, 1Kb, 5Kb, and 10Kb). Correlations between assays were generally consistent across window sizes, both for methylation level and methylation differences (Figure 4). As with gbM, correlations for methylation level were stronger (2-4 fold) than those for methylation differences. Hence, for the coral genome, MBD-seq and mdRAD reproducibly agree with the single-nucleotide measures from WGBS even across small regions.

**Figure 4:**
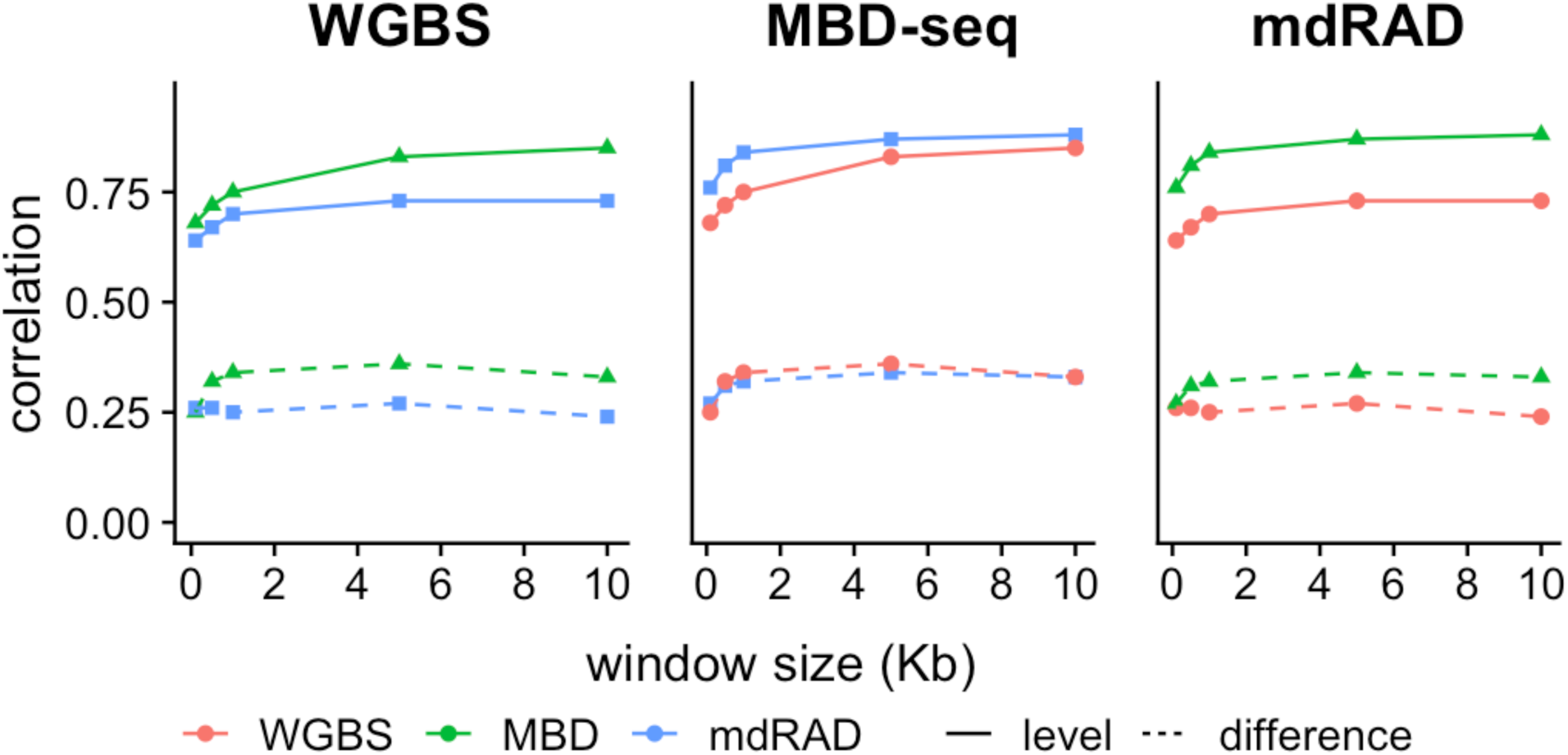
Effect of window size on correlations between assays. Each panel indicates comparisons for one of the assays. Colors indicate the comparison assay. Solid lines indicate correlation of estimates of methylation level for the windows. Dotted lines indicate correlation for estimates of differential methylation between coral colonies.

**Figure 5:**
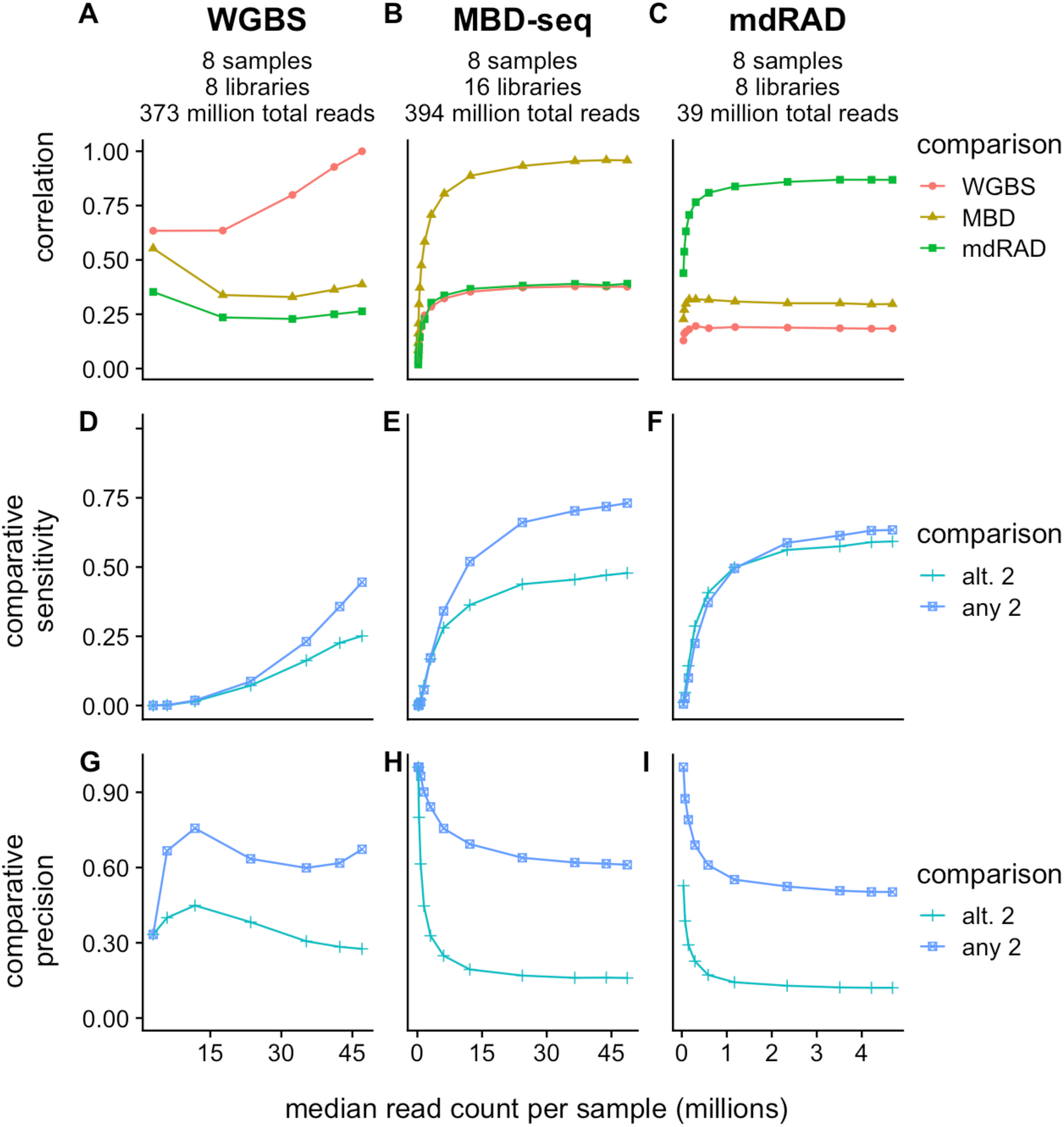
Effect of simulated read reductions on estimates of methylation differences between coral colonies. Columns are assigned to the three assays. Rows are assigned to statistics measuring agreement between assays. Each data point represents a simulated reduction in fold coverage. (A-C) Pearson correlation between assays as fold coverage is reduced. (D-F) Sensitivity of each assay in detecting significant differences (FDR < 0.1) detected by other assays. For each reduction in fold coverage, comparative sensitivity is computed as the number of significant genes shared with the comparison divided the total significant genes for the comparison. Comparisons include *any 2*: genes that were significant in any 2 assays; *alt. 2*: genes that were significant for both the alternative assays (G-I) Precision of each assay in detecting only significant differences (FDR < 0.1) also detected by other assays. For each reduction in fold coverage, comparative precision is computed as the number of significant genes shared with the comparison divided the total significant genes for the fold reduction. Read counts on the X axis refer to the total number of reads included in the final filtered alignment file, hence mapping efficiencies and PCR duplication rates should be accounted for when deciding on total sequencing effort.

### Effect of fold coverage on detecting methylation differences

Given the importance of reducing sequencing costs for ecological epigenetics, we sought to evaluate the importance of sequencing effort for each assay in estimating methylation statistics. To do this, we simulated reduced sequencing effort by random resampling of fold coverage from the datasets. We then re-calculated estimates of methylation level and methylation differences from the reduced sets. As we detected no reproducible differences between polyp types (Figure S12), we focused on differences between colonies (genotype).

For estimates of absolute levels of gbM, fold coverage appeared to matter very little. We found that correlation between assays plateaued between 0.75 and 0.80 with roughly 20% of the original sequencing effort (Figure S13). Although lower, correlation of gbM differences also plateaued with relatively little sequencing effort (Figure 4A-C). Hence correlation between assays was sensitive only to severe reductions in fold coverage. Moreover, increasing fold coverage appeared unlikely to improve correlations between assays.

Detecting significant DMGs in contrast, was more dependent on fold coverage. For the sake of comparability, here we reduced the mdRAD dataset to just eight libraries prepared with the FspE1 enzyme. To illustrate the importance of fold coverage for statistical significance, we plotted the proportion of DMGs detected by at least two of the assays (all overlapping regions in figure 3) that were also detected with each read reduction (‘any 2’ trace in Figure 4 D-F). Given its similarity to the sensitivity metric used to evaluate classification models, we refer to this statistic as comparative sensitivity. For a more stringent test of sensitivity, we also computed this value based on DMGs detected in each of the alternative assays (‘alt. 2’ in Figure 4 D-F). Based on this analysis, it appeared that increasing sequencing effort would have returned many more DMGs for WGBS, somewhat more for MBD-seq and relatively few more for mdRAD. We also assessed how often DMG calls by each assay were corroborated by the other assays. Here we computed comparative precision as the proportion of DMGs from a given reduction that were also significant for at least two of the assays’ full datasets (‘any 2’ in Figure 4G-I). For greater stringency, this was also computed based on significance in the two alternative assays. Corroboration rates were slightly higher for WGBS DMGs, but generally similar for all three assays. When we repeated the analysis using the full mdRAD dataset (which still used less overall sequencing; Table1), mdRAD detected many more corroborated differences, with only slightly lower comparative precision (Figure S14). In summary, mdRAD can identify reproducible differences in methylation with sensitivity and precision comparable to MBD-seq and WGBS with relatively little fold coverage.

## Discussion

Here we present a benchmarking study of methods for assaying DNA methylation for ecological epigenetics in a marine invertebrate. We found that all three assays measure methylation level consistently, with a minimum correlation of 0.8 for gbM (Figure 1). Analysis of differential methylation was less consistent, but still indicated reproducible differences between coral colonies (Figure 2). Surprisingly, we found no such reproducible differences between polyp types (branch tips compared to branch sides; Figure S12). It is interesting to note that in this case WGBS identified 169 DMGs, none of which were detected by the other assays. This may reflect greater sensitivity of WGBS, however, since the other assays identified more of the reproducible differences between genotypes (Figure 2; Figure S13D-F), greater sensitivity seems unlikely. Given the extensive transcriptional differences between axial and radial polyps (Hemond et al. 2014), the absence of reproducible methylation differences between them suggests that variation in gbM is not involved for tissue-specific gene regulation in corals. This result adds to growing evidence that gbM does not directly regulate gene expression in invertebrates (Zilberman 2017; Bewick et al. 2018; Harris et al. 2019).

Simulating reduced sequencing effort for each assay showed that fold coverage is most important in the context of statistical significance. While the number of corroborated DMGs dropped steeply with fold coverage (Figure 4 D-F), correlations between assays were relatively stable (Figure 4 A-C; Figure S8). This suggests that adding a second assay to a methylomic experiment can provide valuable corroboration even with relatively little sequencing effort. This approach could also potentially prevent spurious conclusions. For instance, here we detected no reproducible differences in gbM between polyp types, a conclusion distinct from the one we would have drawn from WGBS alone (over 150 DMGs). Based on these results, we suggest an experimental strategy that uses high fold coverage for one assay to obtain statistical significance and low coverage from one or more other assays for corroboration. For instance, mdRAD could be used to sequence a large number of individuals to identify significant differences, with WGBS, MBD-seq, or both applied with relatively lower coverage for confirmation.

To conclude, MBD-seq and mdRAD are cost effective alternatives to WGBS, providing consistent estimates of methylation level and similar or greater sensitivity to methylation differences at lower library preparation and sequencing costs. The considerably lower sequencing effort required for mdRAD makes it particularly promising for the large sample sizes needed for ecological epigenetics.

## Acknowledgements

This study was supported by the National Science Foundation grant IOS-1755277 to M.V.M. Data analysis was performed with the help of the Texas Advanced Computing Center.

## Data Accessibility

Reads generated for this study have been uploaded to the SRA database project accession PRJNA601565. All scripts for data processing and analysis, as well as intermediate datasets are available on Github (Dixon 2020).

## Notes

https://github.com/grovesdixon/benchmarking_coral_methylation

## References

Akalin, A., Kormaksson, M., Li, S., Garrett-Bakelman, F. E., Figueroa, M. E., Melnick, A., & Mason, C. E. (2012). MethylKit: a comprehensive R package for the analysis of genome-wide DNA methylation profiles. Genome Biology, 13(10). doi: 10.1186/gb-2012-13-10-R87

Andrews, K. R., Good, J. M., Miller, M. R., Luikart, G., & Hohenlohe, P. A. (2016). Harnessing the power of RADseq for ecological and evolutionary genomics. Nat Rev Genet, advance on (2), 81–92. doi: 10.1038/nrg.2015.28

Benjamini, Y., & Hochberg, Y. (1995). Controlling the False Discovery Rate: a Practical and Powerful Approach to Multiple Testing. Journal of the Royal Statistical Society, 57(1), 289–300.

Bewick, A. J., Ji, L., Niederhuth, C. E., Willing, E.-M., Hofmeister, B. T., Shi, X., … Schmitz, R. J. (2016). On the origin and evolutionary consequences of gene body DNA methylation. Proceedings of the National Academy of Sciences of the United States of America, 113(32), 9111–9116. doi: 10.1073/pnas.1604666113

Bewick, A. J., Sanchez, Z., Mckinney, E. C., Moore, A. J., Moore, P. J., & Schmitz, R. J. (2018). Gene-regulatory independent functions for insect DNA methylation. BioRxiv Preprint. doi: https://doi.org/10.1101/355669

Bewick, A. J., Zhang, Y., Wendte, J. M., Zhang, X., & Schmitz, R. J. (2019). Evolutionary and Experimental Loss of Gene Body Methylation and Its Consequence to Gene Expression. 9(August), 2441–2445. doi: 10.1534/g3.119.400365

Bossdorf, O., Richards, C. L., & Pigliucci, M. (2008). Epigenetics for ecologists. Ecology Letters, 11(2), 106–115. doi: 10.1111/j.1461-0248.2007.01130.x

Broad Institute. (2019). Picard Toolkit.

Cesar, H. S. J. (2000). Coral reefs: their functions, threats and economic value. In H. S. J. Cesar (Ed.), Collected Essays on the Economics of Coral Reefs. Kalmar, Sweden: CORDIO, Department for Biology and Environmental Sciences, Kalmar University.

Choi, J., Lyons, D. B., Kim, M. Y., Moore, J. D., Choi, J., Lyons, D. B., … Zilberman, D. (2020). DNA Methylation and Histone H1 Jointly Repress Transposable Elements and Aberrant Intragenic DNA Methylation and Histone H1 Jointly Repress Transposable Elements and Aberrant Intragenic Transcripts. Molecular Cell, 1–14. doi: 10.1016/j.molcel.2019.10.011

Cock, P. J. A., Antao, T., Chang, J. T., Chapman, B. A., Cox, C. J., Dalke, A., … De Hoon, M. J. L. (2009). Biopython: Freely available Python tools for computational molecular biology and bioinformatics. Bioinformatics, 25(11), 1422–1423. doi: 10.1093/bioinformatics/btp163

Dias, B. G., & Ressler, K. J. (2014). Parental olfactory experience influences behavior and neural structure in subsequent generations. Nature Neuroscience, 17(1), 89–96. doi: 10.1038/nn.3594

Dimond, J. L., & Roberts, S. B. (2016). Germline DNA methylation in reef corals: patterns and potential roles in response to environmental change. Molecular Ecology, 25, 1895–1904. doi: 10.1111/mec.13414

Dixon, G. B., Davies, S. W., Aglyamova, G. V, Meyer, E., Bay, L. K., & Matz, M. V. (2015). Genomic determinants of coral heat tolerance across latitudes. Science, 348(6242), 1460–1462.

Dixon, G. (2020). benchmarking coral methylation git repository. https://github.com/grovesdixon/benchmarking_coral_methylation

Dixon, G, Bay, L. K., & Matz, M. V. (2014). Bimodal signatures of germline methylation are linked with gene expression plasticity in the coral Acropora millepora. BMC Genomics, 15, 1109. doi: 10.1186/1471-2164-15-1109

Dixon, G, Bay, L. K., & Matz, M. V. (2016). Evolutionary consequences of DNA methylation in a basal metazoan. Molecular Biology and Evolution, 33(9), 2285–2293. doi: 10.1101/043026

Dixon, Groves, Liao, Y., Bay, L. K., & Matz, M. V. (2018). Role of gene body methylation in acclimatization and adaptation in a basal metazoan. Proceedings of the National Academy of Sciences of the United States of America, 115(52), 13342–13346. doi: 10.1073/pnas.1813749115

Eirin-Lopez, J. M., & Putnam, H. M. (2019). Marine Environmental Epigenetics. Annual Review of Marine Science, 11(1), 335–368. doi: 10.1146/annurev-marine-010318-095114

Elshire, R. J., Glaubitz, J. C., Sun, Q., Poland, J. A., Kawamoto, K., Buckler, E. S., & Mitchell, S. E. (2011). A robust, simple genotyping-by-sequencing (GBS) approach for high diversity species. PLoS ONE, 6(5), 1–10. doi: 10.1371/journal.pone.0019379

Feil, R., & Fraga, M. F. (2012). Epigenetics and the environment: emerging patterns and implications. Nature Reviews. Genetics, 13(2), 97–109. doi: 10.1038/nrg3142

Foden, W. B., Butchart, S. H. M., Stuart, S. N., Vié, J.-C., Akçakaya, H. R., Angulo, A., … Mace, G. M. (2013). Identifying the world’s most climate change vulnerable species: a systematic trait-based assessment of all birds, amphibians and corals. PloS One, 8(6), e65427. doi: 10.1371/journal.pone.0065427

Fuller, Z. L., Mocellin, V. J. L., Morris, L., Cantin, N., Sarre, L., Peng, J., … Przeworski, M. (2019). Population genetics of the coral Acropora millepora : Towards a genomic predictor of bleaching. BioRxiv Preprint. Retrieved from https://doi.org/10.1101/867754

Gavery, M. R., & Roberts, S. B. (2013). Predominant intragenic methylation is associated with gene expression characteristics in a bivalve mollusc. PeerJ, 1, e215. doi: 10.7717/peerj.215

Harris, K. D., Lloyd, J. P. B., Domb, K., Zilberman, D., & Zemach, A. (2019). DNA methylation is maintained with high fidelity in the honey bee germline and exhibits global non-functional fluctuations during somatic development. Epigenetics and Chromatin, 12(1), 1–18. doi: 10.1186/s13072-019-0307-4

Harris, R. A., Wang, T., Coarfa, C., Nagarajan, R. P., Hong, C., Downey, S. L., … Costello, J. F. (2010). Comparison of sequencing-based methods to profile DNA methylation and identification of monoallelic epigenetic modifications. Nature Biotechnology, 28(10), 1097–1105. doi: 10.1038/nbt.1682

Heijmans, B. T., Tobi, E. W., Stein, A. D., Putter, H., Blauw, G. J., Susser, E. S., … Lumey, L. H. (2008). Persistent epigenetic differences associated with prenatal exposure to famine in humans. Proceedings of the National Academy of Sciences of the United States of America, 105(44), 17046–17049. doi: 10.1073/pnas.0806560105

Hofmann, G. E. (2017). Ecological Epigenetics in Marine Metazoans. Frontiers in Marine Science, 4(January), 1–7. doi: 10.3389/fmars.2017.00004

Horsthemke, B. (2018). A critical view on transgenerational epigenetic inheritance in humans. Nature Communications, 9(1), 1–4. doi: 10.1038/s41467-018-05445-5

Irmler, M., Kaspar, D., Hrab, M., Angelis, D., & Beckers, J. (2020). The (not so) Controversial Role of DNA Methylation in Epigenetic Inheritance Across Generations ethylation of Cytosine Represses Gene Expression. In R. Teperino (Ed.), Beyond Our Genes (pp. 175–208). doi: https://doi.org/10.1007/978-3-030-35213-4_10

Kaati, G., Bygren, L. O., Pembrey, M., & Sjöström, M. (2007). Transgenerational response to nutrition, early life circumstances and longevity. European Journal of Human Genetics, 15(7), 784–790. doi: 10.1038/sj.ejhg.5201832

Krueger, F., & Andrews, S. R. (2011). Bismark: A flexible aligner and methylation caller for Bisulfite-Seq applications. Bioinformatics, 27(11), 1571–1572. doi: 10.1093/bioinformatics/btr167

Langmead, B., & Salzberg, S. L. (2012). Fast gapped-read alignment with Bowtie 2. Nature Methods, 9(4), 357–359. doi: 10.1038/nmeth.1923

Lawrence, M., Huber, W., Pagès, H., Aboyoun, P., Carlson, M., Gentleman, R., … Carey, V. J. (2013). Software for Computing and Annotating Genomic Ranges. PLoS Computational Biology, 9(8), 1–10. doi: 10.1371/journal.pcbi.1003118

Li, H., Handsaker, B., Wysoker, A., Fennell, T., Ruan, J., Homer, N., … Durbin, R. (2009). The Sequence Alignment/Map format and SAMtools. Bioinformatics, 25(16), 2078–2079. doi: 10.1093/bioinformatics/btp352

Love, M. I., Huber, W., & Anders, S. (2014). Moderated estimation of fold change and dispersion for RNA-Seq data with DESeq2. Genome Biology, 15(550), 1–21. doi: 10.1101/002832

Martin, M. (2011). Cutadapt removes adapter sequences from high-throughput sequencing reads. EMBnetjournal, 17(1), 10–12.

Matz, M. V., Treml, E. A., Aglyamova, G. V., & Bay, L. K. (2018). Potential and limits for rapid genetic adaptation to warming in a Great Barrier Reef coral. PLoS Genetics, 14(4), 1–19. doi: 10.1371/journal.pgen.1007220

Matz, M. V. (2019). 2bRAD_denovo git repository.

Painter, R. C., Roseboom, T. J., & Bleker, O. P. (2005). Prenatal exposure to the Dutch famine and disease in later life: An overview. Reproductive Toxicology. doi: 10.1016/j.reprotox.2005.04.005

Putnam, H. M., & Gates, R. D. (2015). Preconditioning in the reef-building coral Pocillopora damicornis and the potential for trans-generational acclimatization in coral larvae under future climate change conditions. Journal of Experimental Biology, 218(15), 2365–2372. doi: 10.1242/jeb.123018

Quinlan, A. R., & Hall, I. M. (2010). BEDTools: A flexible suite of utilities for comparing genomic features. Bioinformatics, 26(6), 841–842. doi: 10.1093/bioinformatics/btq033

Radford, E. J., Ito, M., Shi, H., Corish, J. A., Yamazawa, K., Isganaitis, E., … Ferguson-Smith, A. C. (2014). In utero undernourishment perturbs the adult sperm methylome and intergenerational metabolism. Science, 345(6198). doi: 10.1126/science.1255903

Reusch, T. B. H. (2013). Climate change in the oceans : evolutionary versus phenotypically plastic responses of marine animals and plants. Evolutionary Applications, (ISSN 1752-4571). doi: 10.1111/eva.12109

Sarda, S., Zeng, J., Hunt, B. G., & Yi, S. V. (2012). The evolution of invertebrate gene body methylation. Molecular Biology and Evolution, 29(8), 1907–1916. doi: 10.1093/molbev/mss062

Serre, D., Lee, B. H., & Ting, A. H. (2009). MBD-isolated genome sequencing provides a high-throughput and comprehensive survey of DNA methylation in the human genome. Nucleic Acids Research, 38(2), 391–399. doi: 10.1093/nar/gkp992

Strader, M. E., Wong, J. M., Kozal, L. C., Leach, T. S., & Hofmann, G. E. (2019). Parental environments alter DNA methylation in offspring of the purple sea urchin, Strongylocentrotus purpuratus. Journal of Experimental Marine Biology and Ecology, 517(February), 54–64. doi: 10.1016/j.jembe.2019.03.002

Takuno, S., & Gaut, B. S. (2012). Body-methylated genes in arabidopsis thaliana are functionally important and evolve slowly. Molecular Biology and Evolution, 29(1), 219–227. doi: 10.1093/molbev/msr188

Takuno, S., & Gaut, B. S. (2013). Gene body methylation is conserved between plant orthologs and is of evolutionary consequence. Proceedings of the National Academy of Sciences of the United States of America, 110(5), 1797–1802. doi: 10.1073/pnas.1215380110

Takuno, S., Ran, J.-H., & Gaut, B. S. (2016). Evolutionary patterns of genic DNA methylation vary across land plants. Nature Plants, 2(January), 15222. doi: 10.1038/nplants.2015.222

van Oppen, M. J. H., Gates, R. D., Blackall, L. L., Cantin, N., Chakravarti, L. J., Chan, W. Y., … Putnam, H. M. (2017). Shifting paradigms in restoration of the world’s coral reefs. Global Change Biology, 23(9), 3437–3448. doi: 10.1111/gcb.13647

van Oppen, M. J. H., Oliver, J. K., Putnam, H. M., & Gates, R. D. (2015). Building coral reef resilience through assisted evolution. Proceedings of the National Academy of Sciences, 112(8), 1–7. doi: 10.1073/pnas.1422301112

Wang, S., Lv, J., Zhang, L., Dou, J., Sun, Y., Li, X., … Bao, Z. (2015). MethylRAD: A simple and scalable method for genome-wide DNA methylation profiling using methylation-dependent restriction enzymes. Open Biology, 5(11). doi: 10.1098/rsob.150130

Wong, J. M., Johnson, K. M., Kelly, M. W., & Hofmann, G. E. (2018). Transcriptomics reveal transgenerational effects in purple sea urchin embryos: Adult acclimation to upwelling conditions alters the response of their progeny to differential pCO 2 levels. Molecular Ecology, 27(5), 1120–1137. doi: 10.1111/mec.14503

Zemach, A., & Zilberman, D. (2010). Evolution of Eukaryotic DNA Methylation and the Pursuit of Safer Sex. Current Biology, 20(17), R780–R785. doi: 10.1016/j.cub.2010.07.007

Zilberman, D. (2017). An evolutionary case for functional gene body methylation in plants and animals. Genome Biology, 18(1), 87. doi: 10.1186/s13059-017-1230-2

