## Supplementary Figures for "Benchmarking DNA Methylation Assays for Marine Invertebrates"

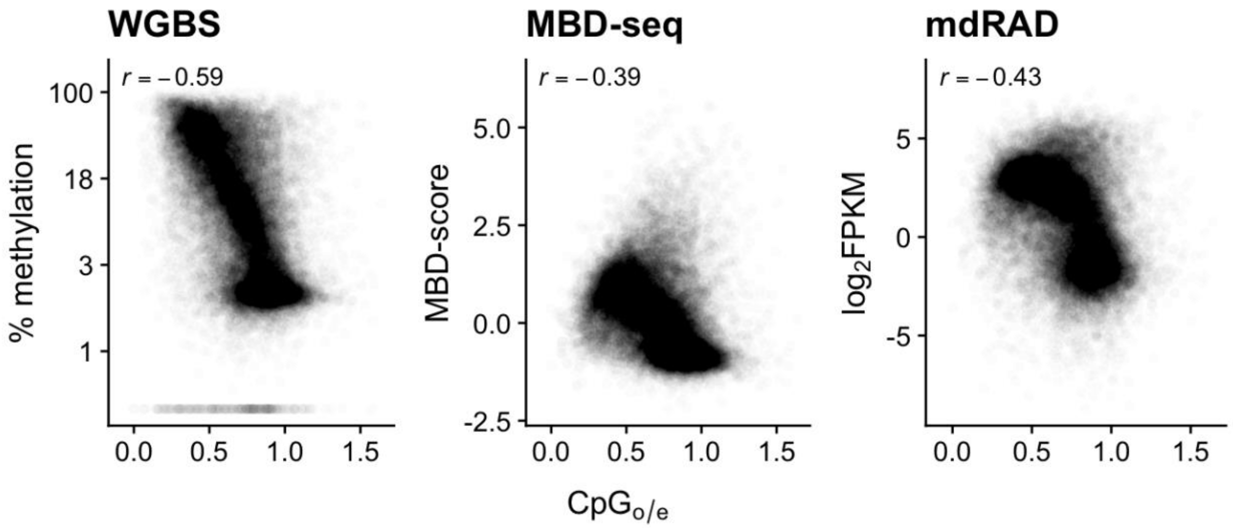


Figure S1: Correlation between gbM level estimates and CpGo/e.

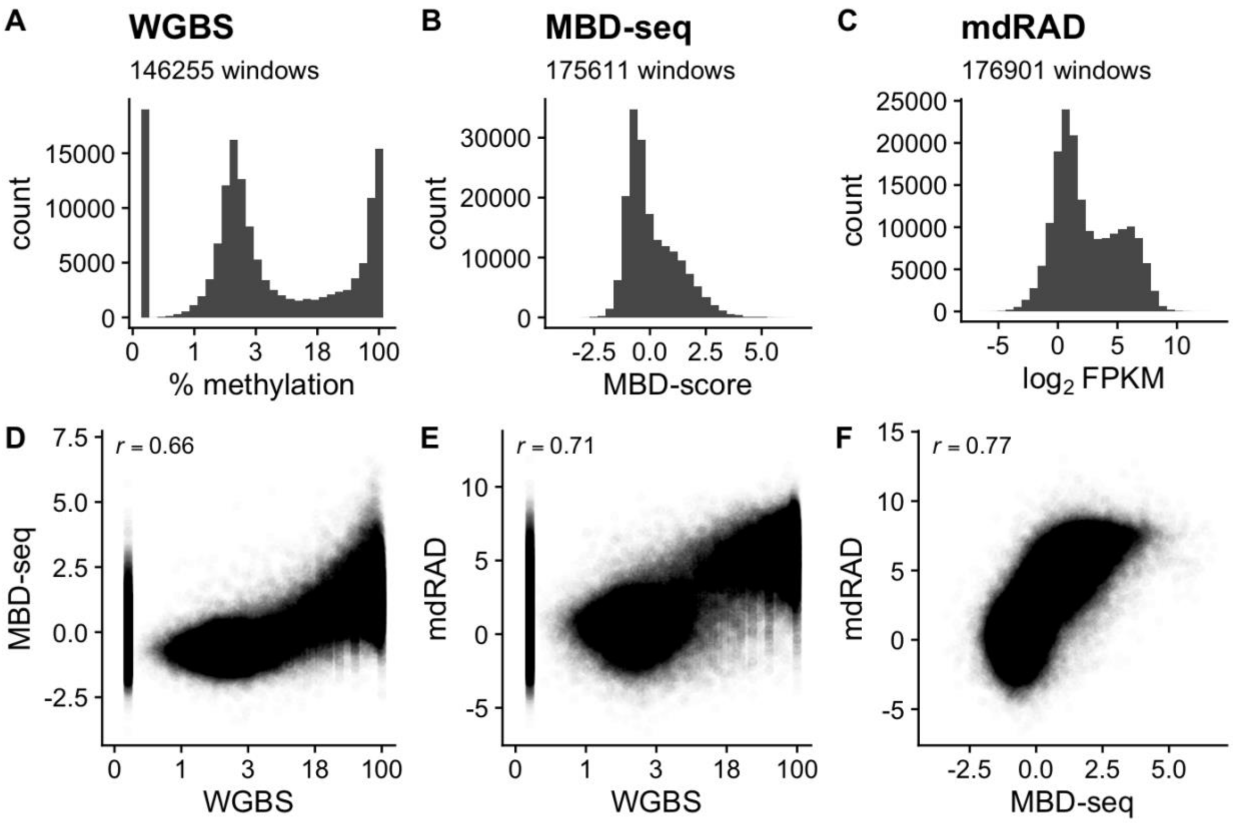


Figure S2: Correlation of methylation level estimates for exons.

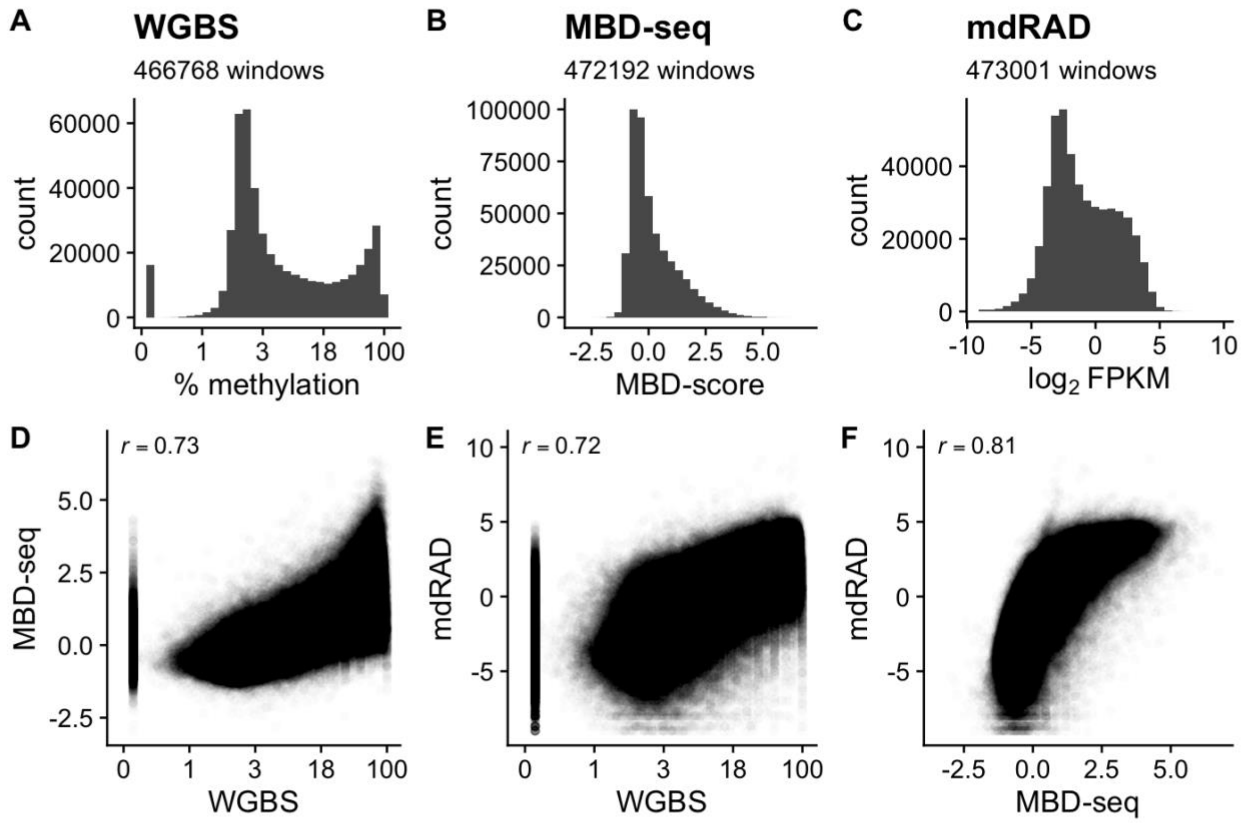


Figure S3: Correlation of methylation level estimates for 1 Kb windows.

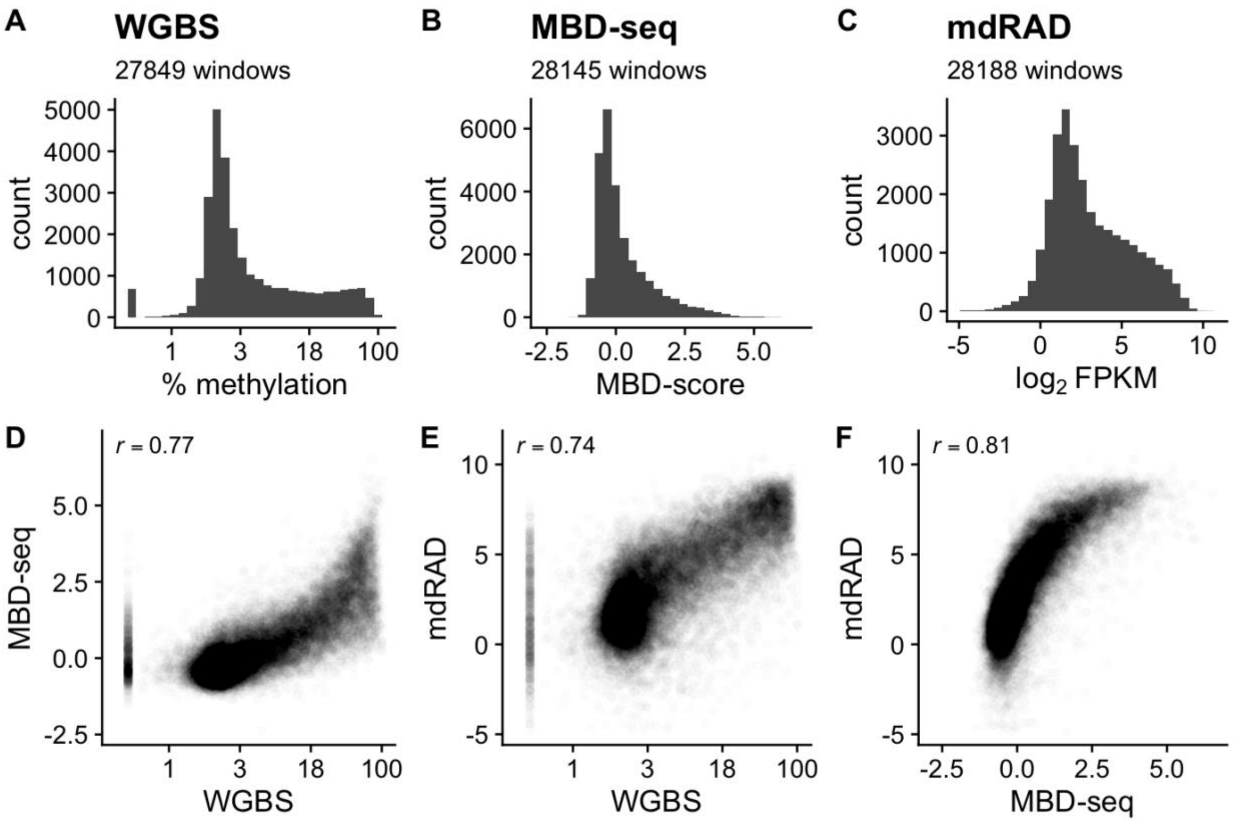


Figure S4: Correlation of methylation level estimates for 1 Kb regions upstream of gene boundaries.

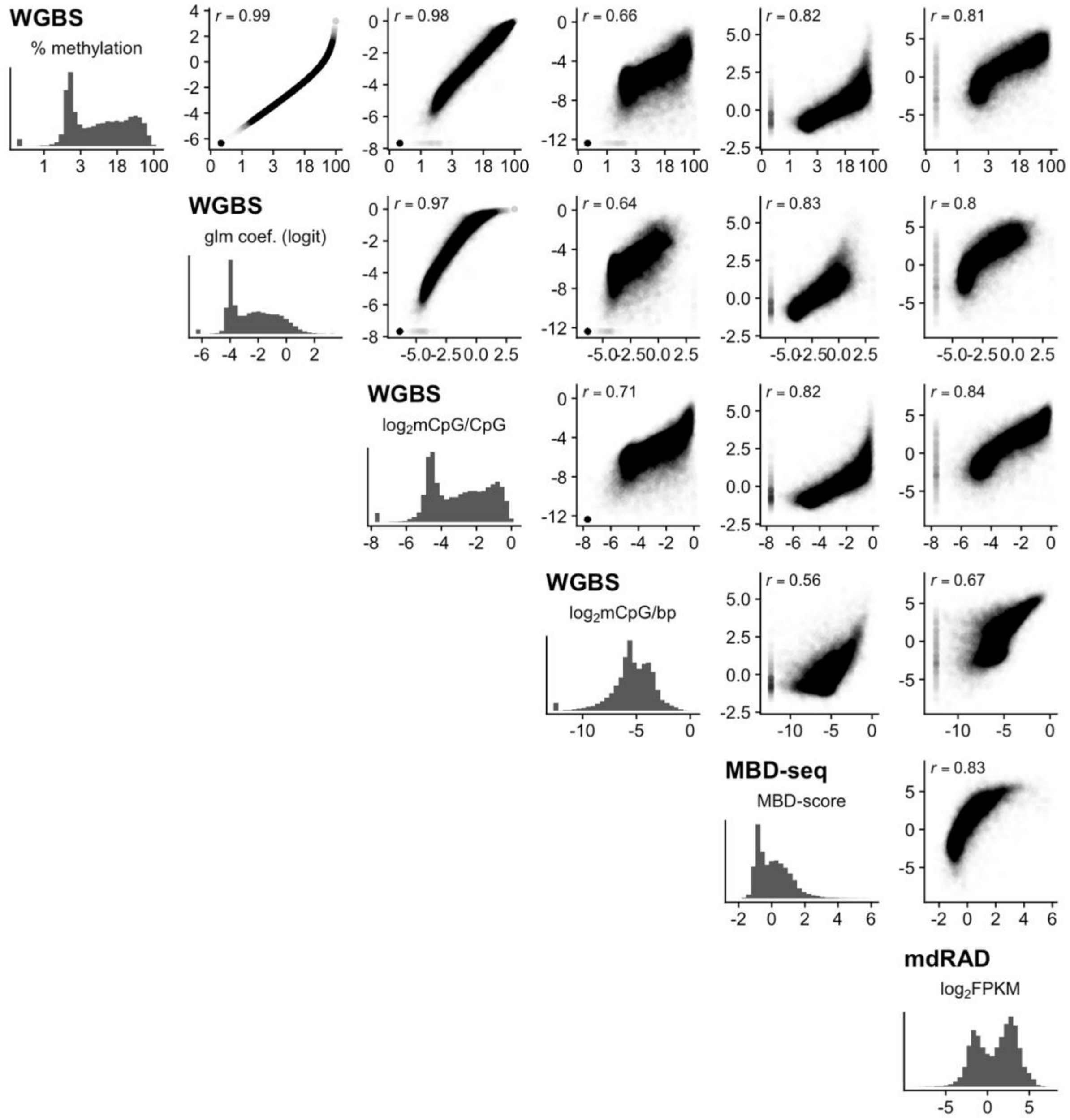


Figure S5: Correlation of gbM level estimates for different metrics based on WGBS. For each panel, the X axis is the metric shown in the histogram on the left side of the row. The Y axis is the metric shown in the histogram at the bottom of the panel's column. Bold titles of the histograms indicate the assay used. The subtitle indicates the particular metric: % methylation is the ratio of methylated fold coverage to total fold coverage summed across all CpG sites in each region and plotted on the log scale; glm coef. (logit) is the sum of the intercept and the coefficient for each region in a generalized logistic regression model of the probability of methylation summed across all CpGs and plotted on the logit scale. $Log_2$mCpG/CpG is the $log_2$ transformed ratio of methylated CpGs to total CpGs. $Log_2$mCpG/bp is the $log_2$ transformed ratio of methylated CpGs to the number of basepairs. MBD-score is the $log_2$ fold difference between the captured fraction and the unbound fraction from each sample's library preparation. $Log_2$FPKM is $log_2$ transformation of the fragments per kilobase of sequence per million reads.

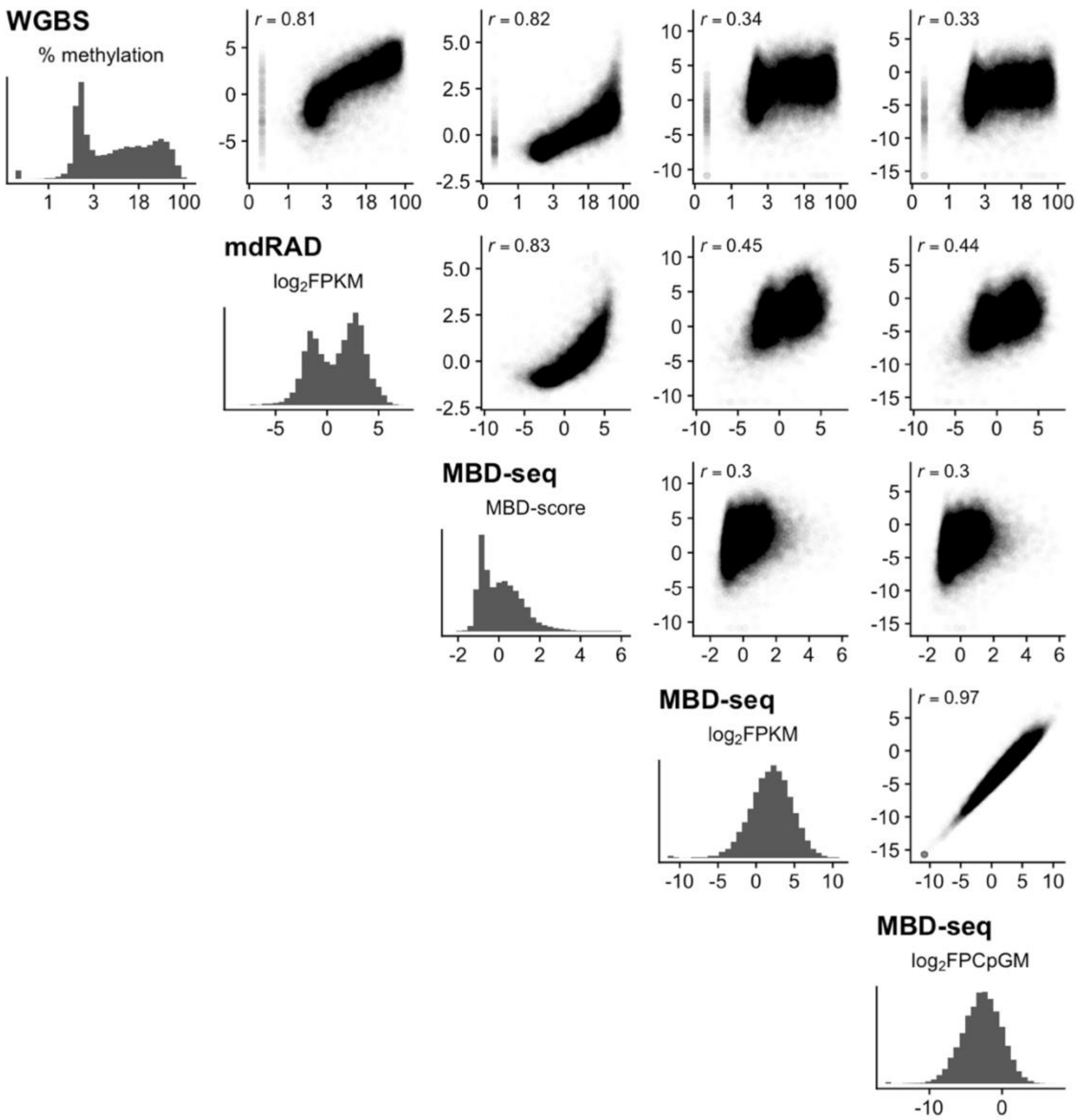


Figure S6: Correlation of gbM level estimates for different metrics based on MBD-seq. For each panel, the X axis is the metric shown in the histogram on the left side of the row. The Y axis is the metric shown in the histogram at the bottom of the panel's column. Bold titles of the histograms indicate the assay used. The subtitle indicates the particular metric: *% methylation* is the ratio of methylated fold coverage to total fold coverage summed across all CpG sites in each region and plotted on the log scale; *glm coef. (logit)* is the sum of the intercept and the coefficient for each region in a generalized logistic regression of the probability of methylation summed across all CpGs plotted on the logit scale. *$Log_2$mCpG/CpG* is the $\log_2$ transformed ratio of methylated CpGs to total CpGs. *$Log_2$mCpG/bp* is the $\log_2$ transformed ratio of methylated CpGs to the number of basepairs. MBD-score is the $\log_2$ fold difference between the captured fraction and the unbound fraction from each sample's library preparation. $Log_2$FPKM is the $\log_2$ transformation of the fragments per kilobase of sequence per million reads. $Log_2$FPCpGM is the $\log_2$ transformation of the fragments per CpG within the region per million reads.

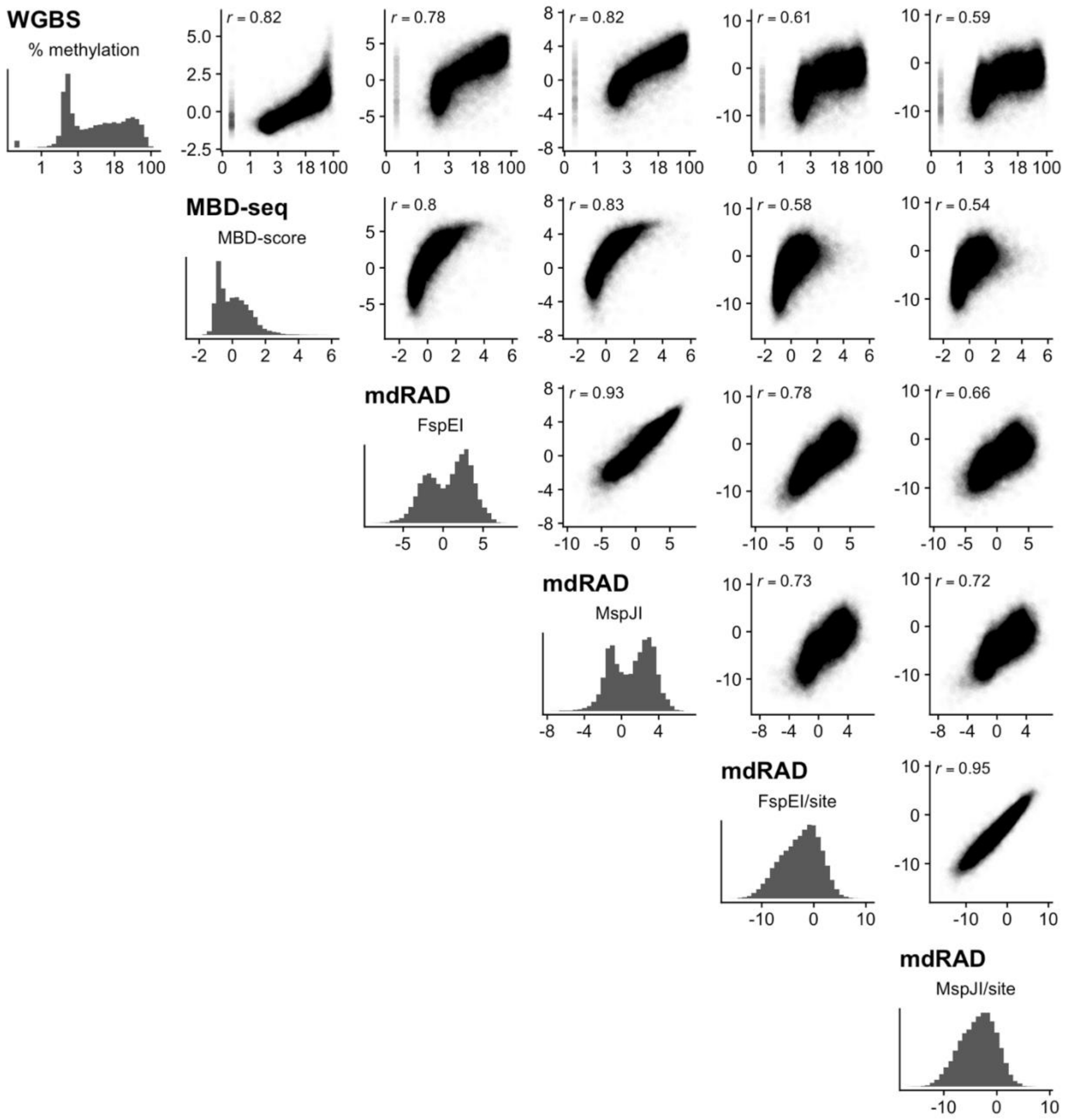


Figure S7: Correlation of gbM level estimates for different metrics based on mdRAD. For each panel, the X axis is the metric shown in the histogram on the left side of the row. The Y axis is the metric shown in the histogram at the bottom of the panel's column. Bold titles of the histograms indicate the assay used. The subtitle indicates the particular metric: *% methylation* is the ratio of methylated fold coverage to total fold coverage summed across all CpG sites in each region and plotted on the log scale; MBD-score is the $\log_2$ fold difference between the captured fraction and the unbound fraction from each sample's library preparation. FspE1 is the $\log_2$ transformation of the fragments per kilobase of sequence per million reads for the FspE1 libraries. MspJ1 is the $\log_2$ transformation of the fragments per kilobase of sequence per million reads for the MspJ1 libraries. The '/site' metrics are $\log_2$ transformation of the fragments per recognition sequence per million reads.

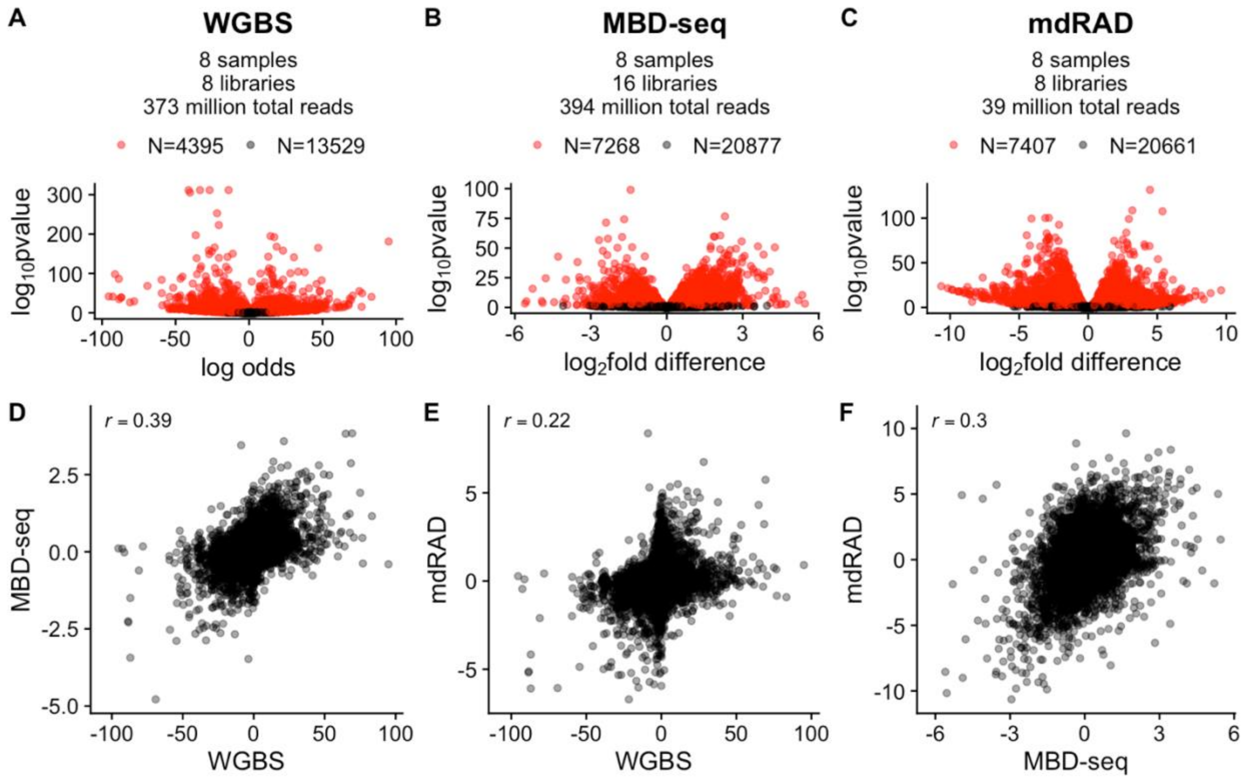


Figure S8: Correlation of gbM difference estimates between two coral colonies (genotypes) using reduced mdRAD dataset with only 8 libraries. (A-C) Volcano plots illustrating differential methylation for the indicated assay. Red points indicate significant regions (FDR < 0.1). The number of biological samples, libraries, final number of aligned reads, and the number of significant and nonsignificant regions is given in the subtitle for each panel. (D-F) Scatterplots of difference estimates. Pearson correlations are indicated in the top left.

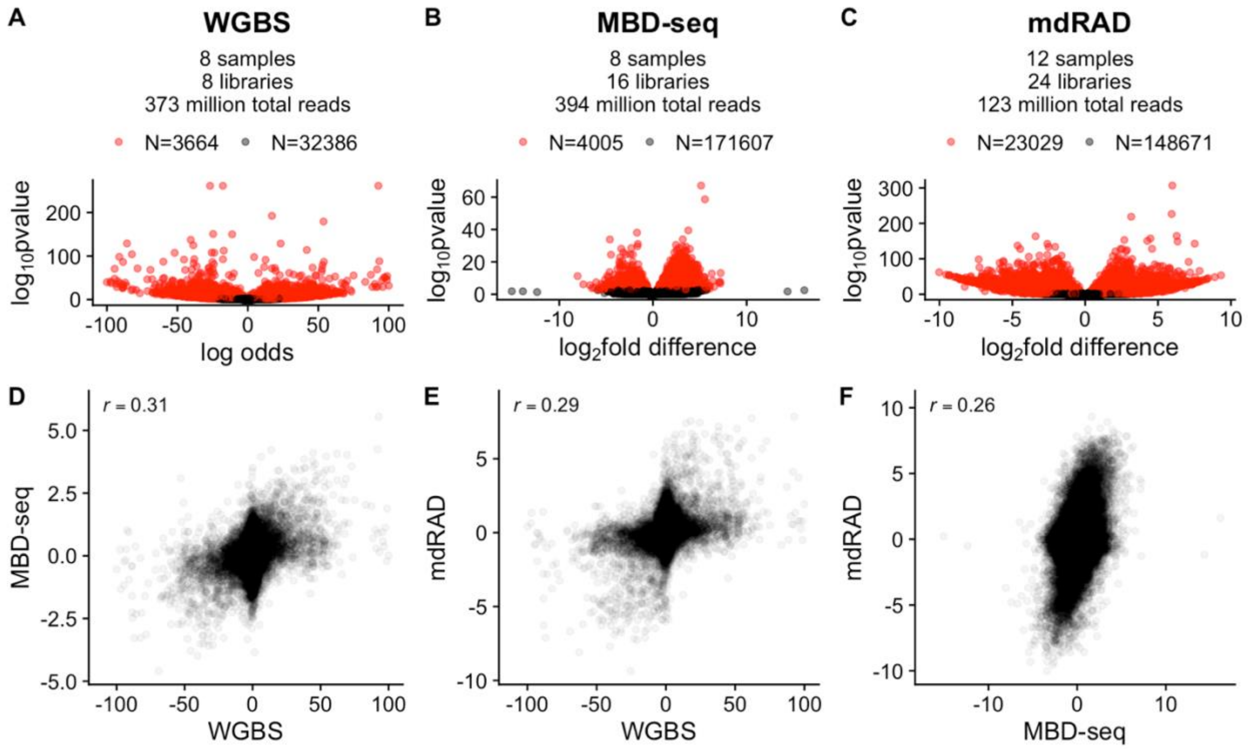


Figure S9: Correlation of methylation difference estimates for exons between two coral colonies (genotypes). (A-C) Volcano plots illustrating differential methylation for the indicated assay. Red points indicate significant regions (FDR < 0.1). The number of biological samples, libraries, final number of aligned reads, and the number of significant and nonsignificant regions is given in the subtitle for each panel. (D-F) Scatterplots of difference estimates. Pearson correlations are indicated in the top left.

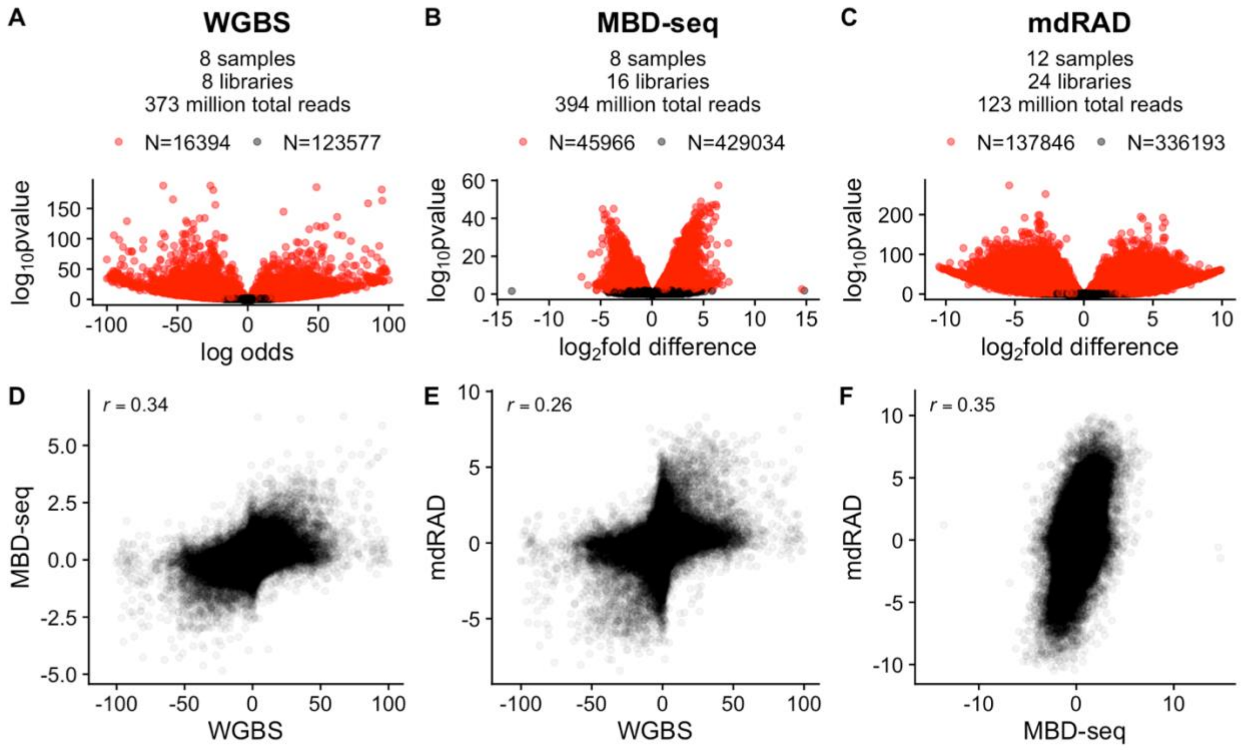


Figure S10: Correlation of methylation difference estimates for 1Kb windows between two coral colonies (genotypes). (A-C) Volcano plots illustrating differential methylation for the indicated assay. Red points indicate significant regions (FDR < 0.1). The number of biological samples, libraries, final number of aligned reads, and the number of significant and nonsignificant regions is given in the subtitle for each panel. (D-F) Scatterplots of difference estimates. Pearson correlations are indicated in the top left.

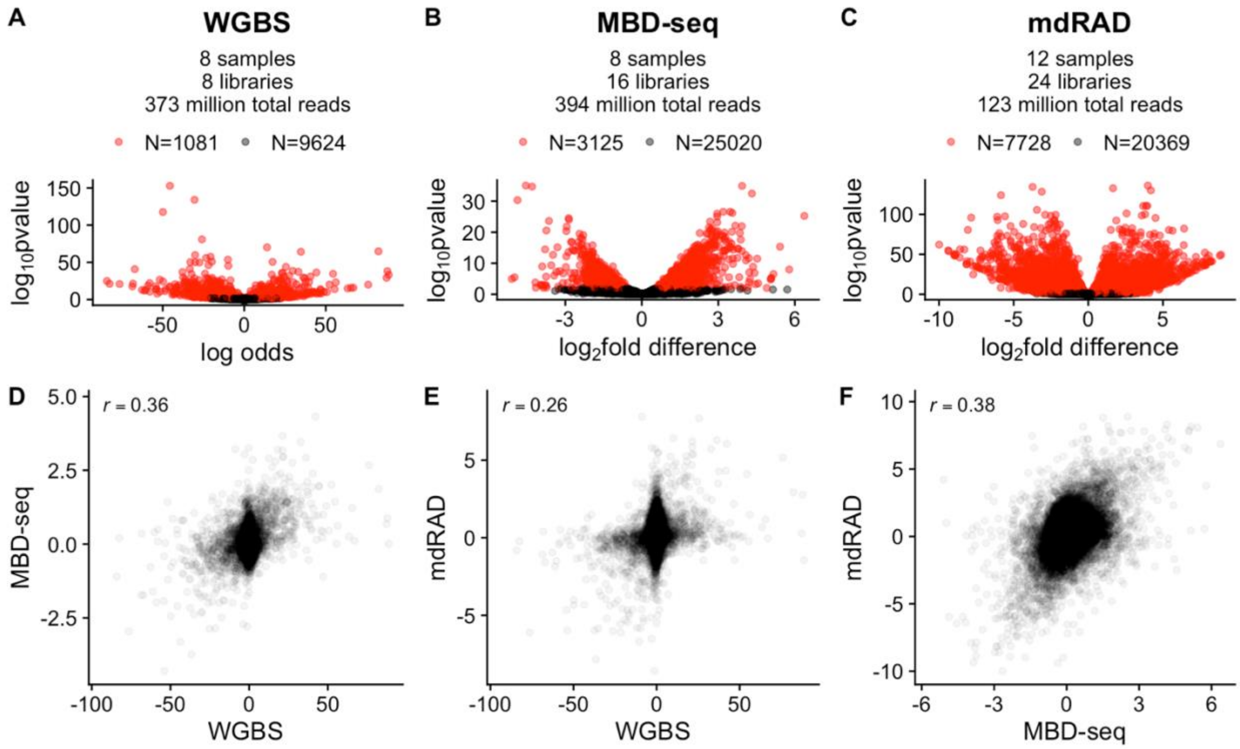


Figure S11: Correlation of methylation difference estimates for upstream regions between two coral colonies (genotypes). (A-C) Volcano plots illustrating differential methylation for the indicated assay. Red points indicate significant regions (FDR < 0.1). The number of biological samples, libraries, final number of aligned reads, and the number of significant and nonsignificant regions is given in the subtitle for each panel. (D-F) Scatterplots of difference estimates. Pearson correlations are indicated in the top left.

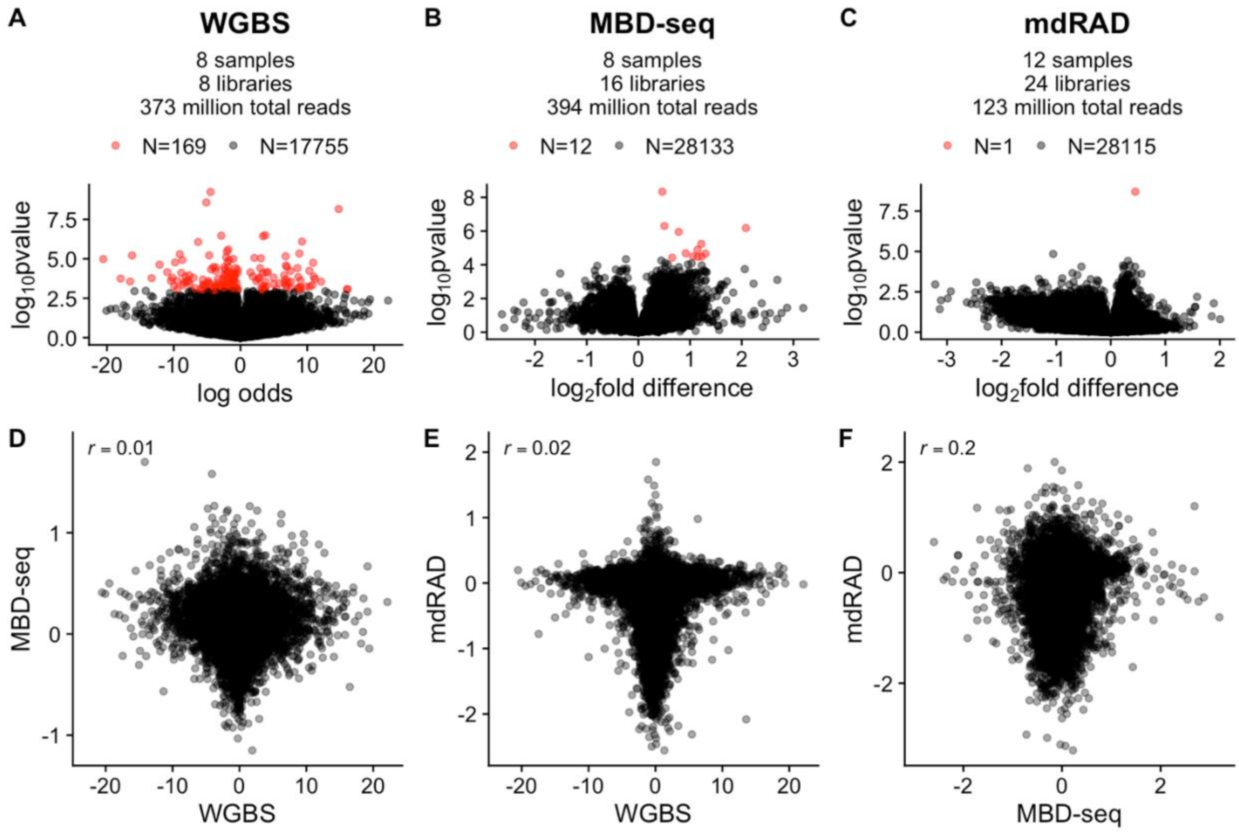


Figure S12: Correlation of gbM difference estimates between branch tips and branch sides. (A-C) Volcano plots illustrating differential methylation for the indicated assay. Red points indicate significant regions (FDR < 0.1). The number of biological samples, libraries, final number of aligned reads, and the number of significant and nonsignificant regions is given in the subtitle for each panel. (D-F) Scatterplots of difference estimates. Pearson correlations are indicated in the top left.

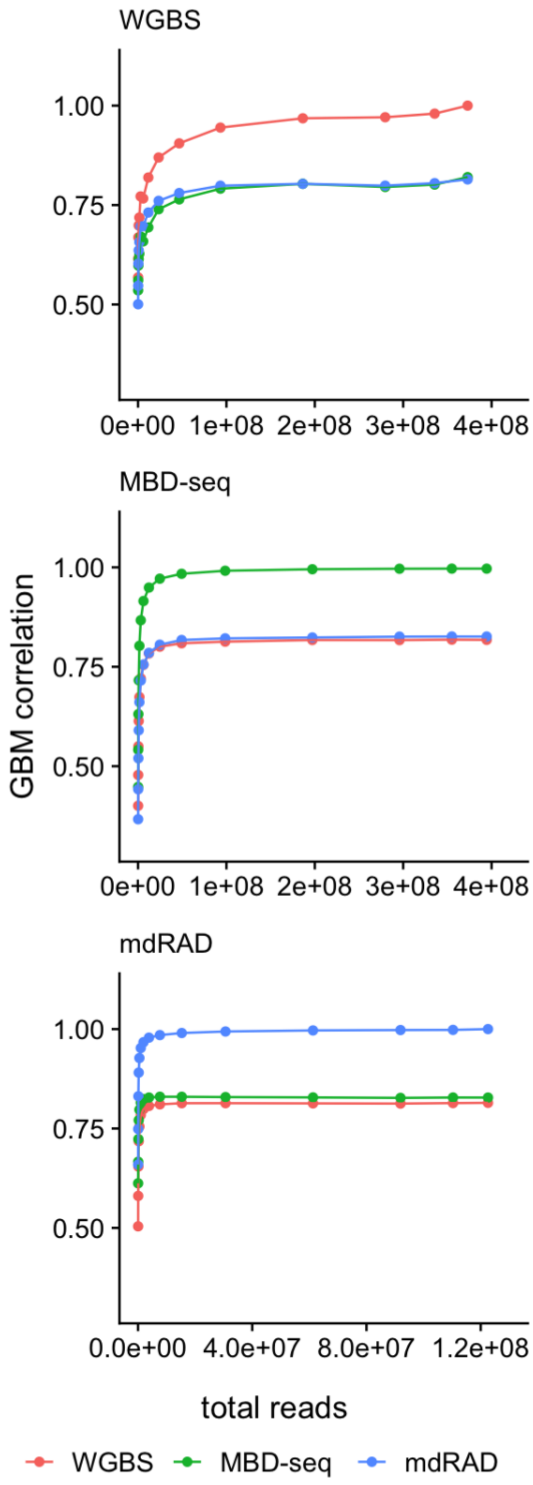


Figure S13: Effect of simulated read reductions on correlation of gbM level estimates from each assay. X axis for each plot shows the simulated total number reads summed across all samples for each assay. The long plateaus seen for each assay show that relatively few reads compared to the amount sequenced are required to maximize concordance on gbM level with other assays.

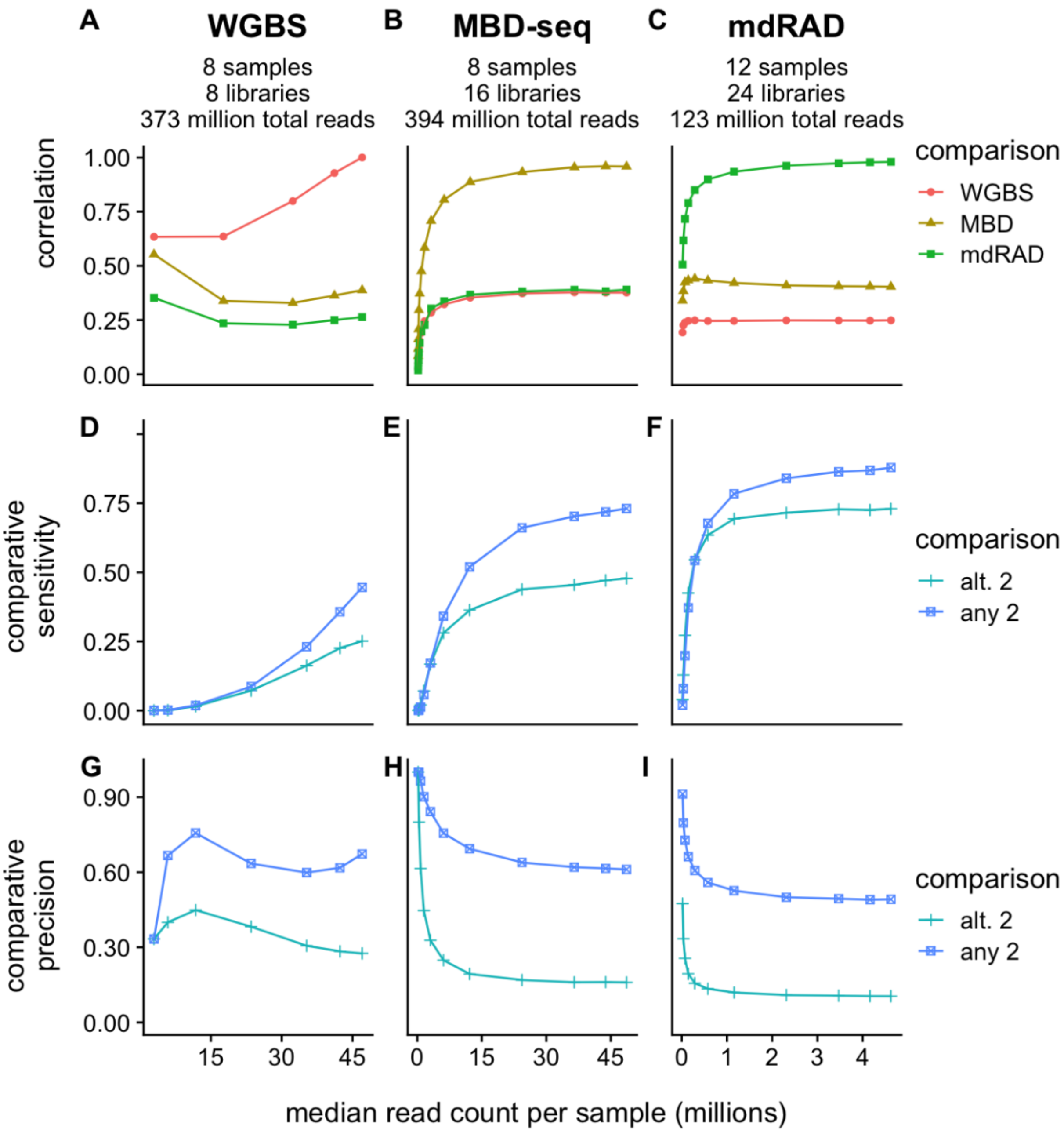

Figure S14: Equivalent to Figure 5 using the full mdRAD dataset.
