## Supplementary Methods for "Benchmarking DNA Methylation Assays for Marine Invertebrates"

Supplementary Methods section for Benchmarking DNA Methylation Assays for Marine Invertebrates

This protocol is for methylGBS, a modified version of methylRAD library preparation (Wang et al. 2015). In the original methylRAD, only ‘double-cut’ fragments (~100bp) were selected and prepared for sequencing. Here instead, we use all fragments with cut sites at either end between 170 and 500 bp in length. This is similar to genotyping by sequencing, but since the enzyme is methylation-selective, the resulting fragments are enriched for methylated DNA. The protocol is similar to the 2bRAD protocol (<https://github.com/z0on/2bRAD_denovo>), but the ligation oligonucleotides are different.

**Digest:**

Digest genomic DNA leaving 4N sticky ends. 100 ng total DNA per digest reaction is suggested input. For DNA isolation procedure see Appendix 1: DNA Isolation.

Digest contents:

| **Reagent** | **amount per reaction** |
| --- | --- |
| Cutsmart buffer | 1x |
| Enzyme activator | 1x |
| FspE1 | 0.4 units total |
| DNA | 100 ng total |

Example Master Mix:

| **Reagent** | **stock concentration** | **vol per reaction (µl)** |
| --- | --- | --- |
| Cutsmart buffer | 10x | 1.5 |
| Enzyme activator | 50x | 0.3 |
| FspE1 | 5 Units / µl | 0.8 |
| water | NA | 10.4 |
| **subtotal** |  | 13 |
| + |  |  |
| DNA | 50 ng/µl | 2.0 |
| **Total** |  | 15.0 |

Steps:

1. Normalize genomic DNA for each sample to a given concentration (eg 50 ng/µl) in water
2. Mix master mix with the appropriate amount of water given the input DNA concentration
3. Aliquot out the master mix into PCR strip tubes or 96 well plate
4. Add DNA to each reaction
5. Set at 37°C for 4 hours
6. Heat kill the enzyme
   a. FspE1: 20 minutes at 80°C
   b. MspJ1: 20 minutes at 65°C
7. Run 4 ul of each digest on gel alongside input to confirm digestion (see next page)
8. Keep on ice and set up ligation same day

**Example digested DNA:**

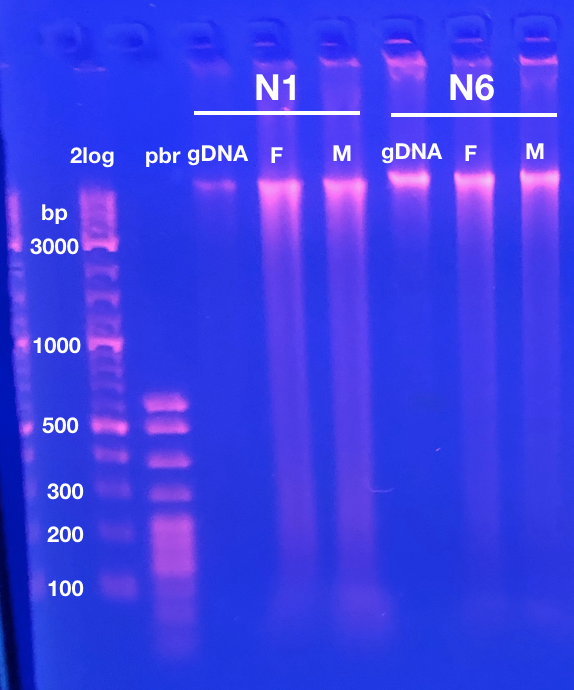

Evidence of digestion was subtle, with additional smear below genomic band, and possibly a slight decrease in size of genomic band.

N1 and N6 were two coral colonies.

2log = 2log ladder

pbr = pBR322 DNA-MspI Digest ladder

gDNA = input DNA

F = FspE1 digest

M = MspJ1 digest

**Ligation:**

Ligate adapters^[1]*^ to the digested DNA.

Ligation contents:

| **Reagent** | **amount per reaction** |
| --- | --- |
| T4 ligase buffer | 1x |
| methylRAD 5ILL adapter | 0.2 µM conc. |
| methylRAD 3ILL adapter | 0.2 µM conc. |
| T4 ligase | 800 units total |
| digest | 10 µl total |

Example Master Mix:

| **Reagent** | **stock concentration** | **vol/rxn (µl)** |
| --- | --- | --- |
| water | na | 4.4 |
| T4 ligase buffer (with 10mM ATP) | 10x | 2.0 |
| 5ILL methylRAD adapter | 5 µM | 0.8 |
| 3ILL methylRAD adapter | 5 µM | 0.8 |
| T4 ligase | 400 units/µl | 2.0 |
| subtotal |  | 10 |
| + |  |  |
| digest | NA | 10 |
| **Total** |  | 20 |

Steps:

1. Make master mix
2. Add 10 ul ligation master mix to each digest (~20 μl final volume)
3. Set at 4°C 6-12 hours (we left ours overnight)
4. Heat kill the ligase with 30 minutes at 65°C

**Amplification:**

Amplify target product using primer binding sites^[2]*^ attached during ligation.

PCR contents:

| **Reagent** | **amount per reaction** |
| --- | --- |
| Titanium taq buffer | 1x |
| dNTPs | 0.3 mM each |
| ILL_Un primer | 0.15 µM conc. |
| ILL-BC primer | 0.15 µM conc. |
| p5 primer | 0.20 µM conc. |
| p7 primer | 0.20 µM conc. |
| Titanium taq | 1x |

Example Master Mix:

| **Reagent** | **stock concentration** | **vol/rxn (µl)** |
| --- | --- | --- |
| water | na | 4.3 |
| Titanium taq buffer | 10x | 2.0 |
| dNTPs | 2.5 mM each | 2.4 |
| ILL_Un primer | 10 µM | 0.3 |
| p5 primer | 10 µM | 0.4 |
| p7 primer | 10 µM | 0.4 |
| Titanium taq | 50x | 0.2 |
| **Subtotal** |  | 10.0 |
| + |  |  |
| ILL-BC primer | 1 µM | 3.0 |
| ligation | na | 7.0 |
| **Total** |  | 20.0 |

Steps:

1. Make master mix without ILL-BC primer or input.
2. Aliquot 10 µl of master mix for each PCR
3. Add appropriate ILL-BC primer and ligation to each reaction
4. Run thermocycler:
   a. 70°C 30 seconds
   b. 16 – 22 cycles of: (95°C 20 seconds; 65°C 3 minutes; 72°C 30 seconds)
   c. 5 minutes at 72°C final extension
5. Run 2 ul on gel (see example below)

**Example amplification gel:**

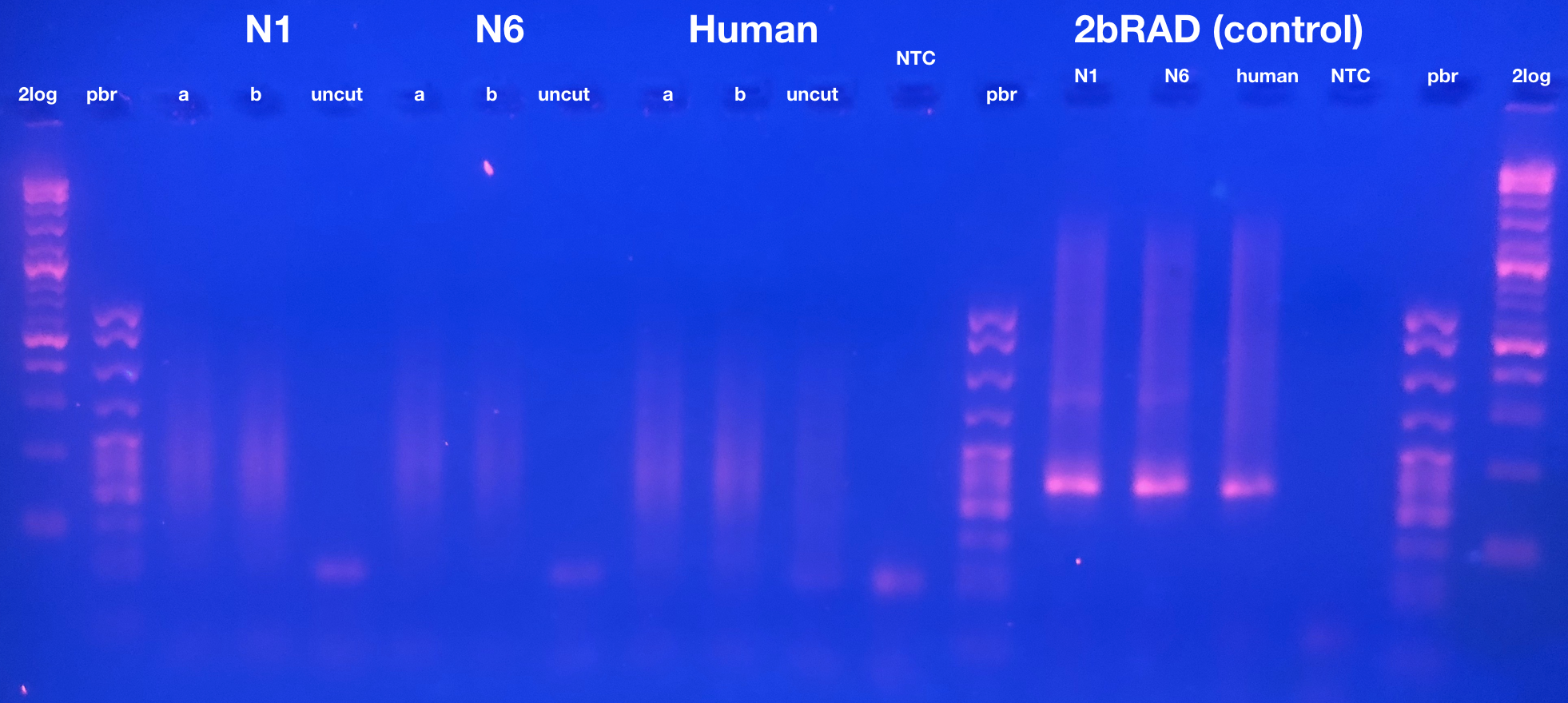

Our amplification did not produce a 100bp band as described in Wang et al. 2015, and selecting this region was not successful. Often smears are even larger in size. Reducing extension time can be used to favor shorter fragments). In the sequencing data, the ‘double-cut’ fragments intended to be selected in Wang et al. 2015 made up only about 1% of the reads.

- N1 and N6 are coral colonies
- ‘a’ and ‘b’ are replicates from each colony
- ‘uncut’ is a negative control using uncut genomic DNA as input for ligation (illustrating that digestion was required to produce PCR product).

**Pool and size select:**

Because samples are now fully barcoded, you can pool them before size selecting.

1. Quantify PCR product from amplification step following picoGreen protocol (see appendix 3 Picogreen assay protocol).
2. Use half of each sample to make a pool with equal amounts of each sample (example table at end of document).
3. Run pool on gel and size select for 170 – 500 bp fragments. The range of sizes can be altered, but Illumina works best with fragments less than 500 bp. It’s advisable to load as much of pool as possible into single well, and not run the gel too long to minimize the total amount of gel you have to purify.
4. Purify the size selected gel fragments with QIAquick Gel Extraction Kit.

**Final test PCR**

Run a final test PCR with P5 and P7 to ensure the library can be amplified

PCR contents:

| **Reagent** | **amount per reaction** |
| --- | --- |
| Titanium taq buffer | 1x |
| dNTPs | 0.3 mM each |
| p5 primer | 0.20 µM conc. |
| p7 primer | 0.20 µM conc. |
| Titanium taq | 1x |

Example Master Mix:

| **Reagent** | **stock concentration** | **vol/rxn (µl)** |
| --- | --- | --- |
| water | na | 13.6 |
| Titanium taq buffer | 10x | 2.0 |
| dNTPs | 2.5 mM each | 2.4 |
| p5 primer | 10 µM | 0.4 |
| p7 primer | 10 µM | 0.4 |
| Titanium taq | 50x | 0.2 |
| **Subtotal** |  | 19.0 |
| + |  |  |
| Size selected pooled library | na | 1.0 |
| **Total** |  | 20.0 |

Steps:

1. Run thermocycler:
   a. 70°C 30 seconds
   b. 12 cycles of: (95°C 20 seconds; 65°C 3 minutes; 72°C 30 seconds)
   c. 5 minutes at 72°C final extension
2. Run 2 ul on gel (see example below)
3. If the gel looks good the library is ready for sequencing.
4. Store the other half of the pool at -20°C as a backup.

**Example final PCR gel:**

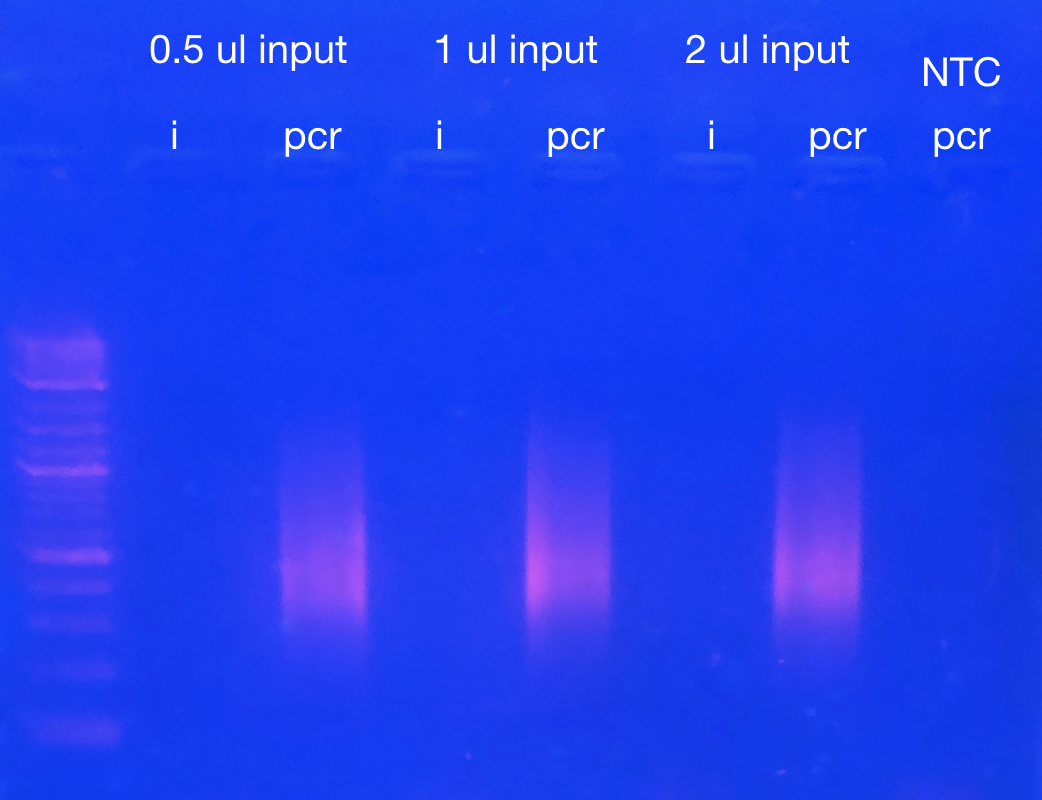

‘i’ indicates equivalent amount of library that was used as input for the PCR. ‘pcr’ indicates product. Note these fragments were selected to be 170 – 800 bp, but we suggested selecting for 170 – 500 because fragments larger than 500 bp do not work well for Illumina sequencing.

Table 1: Example of pooling after Picogreen. Sample numbers are arbitrary. ‘f’ indicates FspE1 digested. ‘m’ indicates MspJ1 digested. ng/µl is the concentration of the amplified/barcoded library based on picogreen. I then calculated the volume needed for 200 ng for each sample. I then added that volume from each sample into a pooled tube. The final concentration and volume of the pool are given below.

| **number** | **sample** | **ng/µl** | **vol for 200 ng** |  | |
| --- | --- | --- | --- | --- | --- |
| 1 | f9 | 44.1 | 4.5 |  | |
| 2 | f10 | 35.2 | 5.7 |  | |
| 3 | f11 | 41.4 | 4.8 |  | |
| 4 | f12 | 40.8 | 4.9 |  | |
| 5 | f13 | 40.2 | 5 |  | |
| 6 | f14 | 37.2 | 5.4 |  | |
| 7 | f15 | 28.3 | 7.1 |  | |
| 8 | f16 | 23.1 | 8.7 |  | |
| 9 | f17 | 36.8 | 5.4 |  | |
| 10 | f18 | 24.6 | 8.1 |  | |
| 11 | f19 | 40.5 | 4.9 |  | |
| 12 | f20 | 30.2 | 6.6 |  | |
| 13 | m9 | 35.6 | 5.6 |  | |
| 14 | m10 | 36.4 | 5.5 |  | |
| 15 | m11 | 19.1 | 10.5 |  | |
| 16 | m12 | 27.3 | 7.3 |  | |
| 17 | m13 | 45.7 | 4.4 |  | |
| 18 | m14 | 37.9 | 5.3 |  | |
| 19 | m15 | 34.9 | 5.7 |  | |
| 20 | m16 | 41.9 | 4.8 |  | |
| 21 | m17 | 25.6 | 7.8 |  | |
| 22 | m18 | 22.1 | 9 |  | |
| 23 | m19 | 22.8 | 8.8 |  | |
| 24 | m20 | 29.2 | 6.8 |  | |
|  |  |  |  | **pool vol.** | **pool conc. (ng/µl)** |
|  |  |  |  | 152.7 | 31.4 |

**Appendix 1: DNA Isolation**

**Reagents:**

1. Lysis buffer (RNAqueous™ Total RNA Isolation Kit (cat no. AM1912). Alternatively use dispersion buffer (see appendix 2: making dispersion buffer)
2. Phenol:Chloroform:Isoamyl Alcohol (25:24:1)(PCA)
3. Sodium Acetate (3 M)(NaOAc)
4. 100% isopropanol
5. 80% ethanol
6. RNAse A (10 mg/mL)(cat no. EN0531)
7. Zymo Clean and concentrator kit (cat no. D4011)

**Steps:**

1. Disrupt tissue in 600 μl of lysis buffer
   - This can be done with a razor in a petri dish or using a bead beater
   - For this study, disrupted 4 axial polyps, or equivalent amount of radial polyps in pooled total of 2 ml lysis buffer, disrupted with a razor in a petri dish, homogenized by pipetting up and down, then split into three separate 2 ml tubes (~600 µl per tube) to continue isolation.
2. Vortex and then incubate lysis buffer and disrupted tissue at 42°C for 2 hours
3. Spin briefly and transfer supernatant to new tube (leaving skeletal debris)
4. Add 600 µl PCA (equal to volume of disrupted tissue and lysis buffer) to lysate, vortex, then chill by placing on ice for 1 minute, then vortex again
5. Spin at 4°C at max speed for 20 minutes
6. Transfer aqueous phase (~550 µl) to new 1.5 ml tube
7. Add 55 µl NaOAc (1/10^th^ the volume) and 363 µl (2/3^rd^ the volume) 100% isopropanol and mix by inverting 20 times
8. Spin at 4°C at max speed for 30 minutes to pellet DNA
9. Pour off supernatant slowly into liquid waste
10. Add 1 ml 80% ethanol and gently wash around tube
11. Pour off 80% ethanol, then briefly spin and pipet away any residual ethanol
12. Resuspend in 50 µl milliQ water
13. Add 1 µl RNAse A and incubate 1 hour at room temperature
14. Clean DNA with Zymo clean and concentrator Kit according to protocol

**Appendix 2: Making dispersion buffer**

*Recipe for dispersion buffer comes from Generation of cDNA Libraries page 106*

*Handle all reagents under fume hood

- Materials: Guanadine thiocyanate (light sensitive, stored in flame cabinet, may be wrapped in foil), sodium citrate dihydrate (Na3C6H5O7 .2H2O; pH of a solution at 25.0°C 7.0-9.0), beta-mercaptoethanol (stored in flame cabinet), milliQ water.
- Equipment: Plate with stirring rod, beaker for mixing, glass bottle for storage.

**Buffer Contents:**

| **Reagent** | **Target Concentration** | **Molecular Mass** |
| --- | --- | --- |
| Guanadine thiocyanate | 4 M | 118.16 |
| Sodium Citrate dihydrate | 30 mM | 294.10 |
| Β-mercaptoethanol | 30 mM | Stock concentration = 14.3 M |

**Recipe for making 50 ml of buffer:**

Guanadine thiocyanate: 23.632 g

Sodium citrate dihydrate: 0.441 g

β-mercaptoethanol: 105 μl

1. Set up stirring plate under hood
2. Set 50 ml of milli Q water stirring
3. Add reagents slowly to stirring liquid
4. transfer dispersion buffer to labeled storage buffer
5. Store at 4°C protected from light

**Appendix 3: Picogreen assay protocol**

1) Place 100ul 1X TE into all first column wells except B1
2) Add 150ul of stock curve (@ 2ug/ml) into B1
3) Serially dilute standards by taking 50ul of B1, mixing into C1, taking 50ul of C1, mixing into D1, and so on until taking 50ul from H1 and throwing it out.
4) To all sample wells, add 98ul of 1X TE.
5) Add 2ul sample DNA to sample wells.
6) Mix Pico Green Master mix: 99.5ul 1XTE + 0.5ul PicoGreen for one sample. Multiply accordingly (plus 8 wells for DNA standard).
7) Add 100ul of master mix to all standard and sample wells, bringing up final volumes in each well to 200.
8) Turn on the plate reader (SpectraMaxM2 in our case).
9) Open software (SoftMaxPro V5 in our case).
10) Select the premade program and run it. Read the fluorescence (excitation 480nm, emission 520nm).
11) Save the data into txt file, assemble the results in Excel in two-column form – well, reading - save it as comma-delimited (.csv) file. The file must contain all A1-H1 wells (blank and calibrators) plus an arbitrary number of sample wells, in any order.
12) Use picogreen.R script to calculate sample concentrations (ng/ul in the original sample).

1. ^*^ Note that while the adapters are similar to 2bRAD their sequences are not the same. Be sure to use the methylRAD oligos.
2. * Note that 2bRAD primers will work here
